# Dengue virus replicative-form dsRNA is recognized by RIG-I and MDA5 cooperatively to activate innate immunity

**DOI:** 10.1101/2024.10.15.618382

**Authors:** Sichao Ye, Yisha Liang, Yu Chang, Bailiang Lai, Jin Zhong

## Abstract

RIG-I like receptors (RLRs) are a family of cytosolic RNA sensors that sense RNA virus infection to activate innate immune response. It is generally believed that different RNA viruses are recognized by either RIG-I or MDA5, two important RLR members, depending on the nature of pathogen-associated molecular patterns (PAMPs) that are generated by RNA virus replication. Dengue virus (DENV) is an important RNA virus causing serious human diseases. Despite extensive investigations, the molecular basis of the DENV PAMP recognized by the host RLR has been poorly defined, and which RLR is involved in sensing the DENV PAMP remains controversial. Here, we demonstrated that the DENV infection-induced interferon response is dependent upon both RIG-I and MDA5, with RIG-I playing a predominant role. Next we purified the DENV PAMP RNA from the DENV-infected cells, and demonstrated that the purified DENV PAMP is viral full-length double-stranded RNA bearing 5’ppp modifications, likely representing the viral replicative-form RNA. Finally, we confirmed the nature of the DENV PAMP by reconstituting the viral replicative-form RNA from *in vitro* synthesized DENV genomic RNA. In conclusion, our work not only defined the molecular basis of the RLR-PAMP interaction during DENV infection, but also revealed the previously underappreciated recognition of the distinct moiety of same PAMP by different RLRs in innate immunity against RNA viruses.

**Importance:** The molecular interaction between PAMPs and RLRs plays a crucial role in innate immune response against the RNA virus infection. To our knowledge, the exact molecular basis of the DENV PAMPs and which RLR member (RIG-I or MDA5) is involved in this recognition remain controversial. In this study, we demonstrated that the DENV PAMP is likely DENV replicative-form RNA, a double-stranded RNA. RIG-I and MDA5 can sense different moieties of this DENV PAMP to active innate immune response. Our work not only clarified which RLR and what viral PAMP are involved in innate immune sensing of DENV infection, but also revealed previously underappreciated recognition of the same PAMPs by different RLRs in innate immunity against RNA viruses.

## Introduction

Dengue virus (DENV) belongs to the family *Flaviviridae*, genus *Flavivirus*, and exists as four antigenically distinct serotypes (denoted DENV1–4)(1). DENV is predominantly transmitted through bites of female mosquitoes *Aedes aegypti* and *Aedes albopictus*, and is recognized as one of the most severe arboviral infections around the world(2). Approximately half of the global population is currently at risk of DENA infection, with about 100 to 400 million infections occurring annually. DENV infection causes a wide spectrum of diseases, ranging from asymptomatic infections to febrile illnesses, and potentially progressing to life-threatening dengue hemorrhagic fever and dengue shock syndrome(3).

DENV is a positive (+) strand RNA virus. The 10.7-kb RNA genome consists of a capped 5’ untranslated region (5’UTR), a single open reading frame (ORF), and a 3’UTR(4). The viral ORF is translated into a polyprotein, which is then processed to individual proteins including three structural proteins (C, precursor membrane (prM), and E) as well as seven non-structural proteins (NS1, NS2A, NS2B, NS3, NS4A, NS4B, and NS5)(5). The DENV entry is achieved through receptor-mediated endocytosis. Upon the fusion of viral envelope and endosomal membrane, the DENV nucleocapsid disassembles to release the viral genomic RNA into the cytoplasm. The viral genomic RNA serves as the mRNA template for the synthesis of viral proteins. The viral RNA-dependent RNA polymerase (NS5) replicates the viral RNA genome within replication complexes formed on the endoplasmic reticulum membrane(6). Throughout this intricate process, multiple forms of viral RNA are generated. The genomic (+) sense RNA is used as a template to synthesize a negative (−) sense RNA, which remains base-paired with the (+) sense RNA to form a double-stranded RNA (dsRNA) called replicative form (RF). Subsequently, the (−) sense RNA within the RF acts as a template for the synthesis of nascent (+) sense RNA. As the nascent (+) sense RNA elongates, it displaces the original (+) sense RNA and remains base pairing with the (−) sense RNA template, forming a three-strand RNA complex called replicative intermediate (RI)(4).

The host innate immune system, acting as the primary defense against invading pathogens, employs a variety of pattern recognition receptors (PRRs) to recognize conserved molecular structures of pathogens, known as pathogen-associated molecular patterns (PAMPs), which are often vital for the pathogen life cycle(7). Retinoic acid-inducible gene I (RIG-I)-like receptors (RLRs), a family of cytosolic PRRs that recognize RNA structure-based PAMPs, comprise three members: RIG-1, melanoma differentiation-associated protein 5 (MDA5) and laboratory of genetics and physiology 2 (LGP2)(8). All RLRs contain a central helicase domain and a carboxy-terminal domain (CTD), which collaboratively detect RNA ligands. Unlike LGP2, RIG-I and MDA5 contain two additional N-terminal caspase activation and recruitment domain (CARD), which is responsible for subsequent signal transduction. As the result, RIG-I and MDA5 can directly recognize RNA ligands and transduce the signals, whereas LGP2 lacks CARD and is believed to play a regulatory role in RIG-I and MDA5 sensing(9, 10). It has been shown that RIG-I and MDA5 recognize RNA ligands with distinct molecular features. RIG-I detects short dsRNA with a 5’-triphosphate (5’ppp) end(11–14). In addition, RIG-I can detect 5’ppp-single stranded RNA (ssRNA) with certain molecular features yet to be determined. For example, the polyU/UC tract within the hepatitis C virus (HCV) 3’UTR has been demonstrated to activate RIG-I(15). On the contrary, MDA5 detects long dsRNA(16). When RIG-I or MDA5 binds to RNA ligands, they undergo conformational changes that facilitate their interaction with CARD domain located in the critical adaptor protein mitochondrial antiviral signaling protein (MAVS). This intermolecular CARD-CARD interaction initiates a signaling cascade that activates the transcription factors NF-κB, interferon regulatory factor 3 (IRF3) and IRF7, which in turn triggers the transcription of the proinflammatory cytokines and the expression of type I and III interferons (IFNs)(10, 17).

Previous studies showed that MAVS deficiency strongly impairs the induction of cytokines and IFNs in response to DENV infection both in *vivo*(18) and *in vitro*(19–21), supporting an important role of the RLR-mediated signaling in innate immune response to DENV infection. However, the molecular identity of DENV PAMPs recognized by RLRs has been poorly defined. Maxime and colleagues(22) performed protein affinity purification by overexpressing RIG-I in DENV-infected HEK293 cells, and found that RIG-I-bound DENV RNAs are mainly single-stranded (+) sense viral genomic RNAs with 5’ppp. Therefore, they proposed that the RIG-I recognizes certain secondary structures located within the 5’region of the nascent uncapped single-stranded DENV genomic RNA. However, this experiment was performed in cells overexpressing RLRs, and it is imperative to identify the exact viral RNA species recognized by RLRs under physiological conditions. In addition, it remains controversial which RLR is involved in sensing DENV PAMPs to active innate immune response. It is reported that DENV-induced IFN responses in African green monkey kidney Vero cells or human lung carcinoma A549 cells were abolished by overexpressing a dominant-negative RIG-I(23), and the IFN response was significantly diminished in DENV-infected human brain microvascular endothelial cells following RIG-I knockdown(24), whereas the knockdown of MDA5 in HEK293 cells had no effect(22). These results indicate an important role of RIG-I rather than MDA5 in mediating IFN response to the DENV infection. However, it was shown that the IFN response in DENV-infected mouse fibroblast cells was only blocked by silencing both RIG-I and MDA5 but not by silencing either of them(19). Consistently, the DENV replication was only enhanced in A549 cells where both RIG-I and MDA5 were silenced while remained unchanged in the RIG-I- or MDA5-knockdown cells (25). In addition, other studies reported that the silencing RIG-I or MDA5 alone could reduce the IFN response upon DENV infection in human dendritic cells(20, 21). These results indicate that both RIG-I and MDA5 may be involved in mediating the innate immune response against DENV infection.

In this study, we found that both RIG-I and MDA5 were involved in the innate sensing of DENV infection, with RIG-I playing a main role. Furthermore, we demonstrated that RIG-I and MDA5 cooperatively recognized a different moiety in the DENV RF dsRNA to activate the innate immune response.

## Results

### DENV2 infection-induced IFN response is dependent upon both MDA5 and RIG-I, with RIG-I playing a predominant role

DENV employs a multitude of strategies to evade the host innate immune responses, which involves multiple viral proteins such as NS2A, NS2B, NS3, NS4A, NS4B, and NS5(26–31), thus we first evaluated the induction of IFN response in different DENV-infected cell lines. We infected HEK293 cells with DENV2 strain 16681 at a multiplicity of infection (MOI) of 1. Type I and III IFNs as well as intracellular DENV2 RNA levels were determined at the indicated time points post-infection by RT-qPCR. DENV2 infection elicited modest interferon response at 24 hours post-infection, but the IFN production dramatically increased at 48 hours post-infection (Fig. 1A-1B). The increased IFN response was found to correlate with the replication of DENV2 RNA (Fig. 1C). Similar to HEK293 cells, DENV2 infection induced robust IFN response in Huh7 and Vero cells (Fig. S1). These results suggested that despite DENV is able to antagonize the host innate immunity to a certain degree, it still triggers strong IFN response in a variety of cells.

**Fig. 1.**
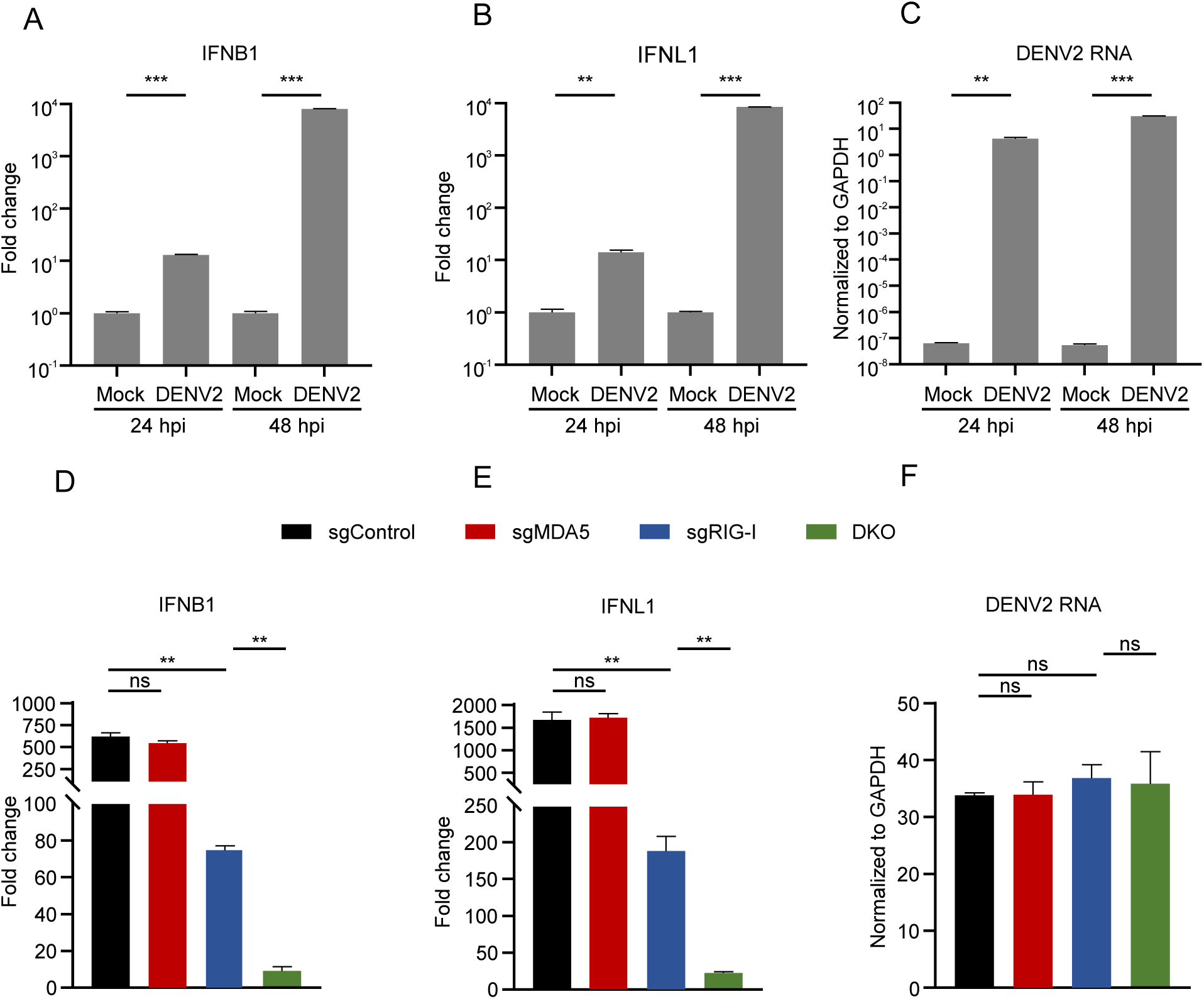
DENV2 infection-induced IFN response is dependent upon both MDA5 and RIG-I, with RIG-I playing a predominant role. (**A-C**) HEK293 cells were infected with DENV2 (MOI=1). Infected cells were analyzed at indicated time points for IFNB1 (**A**), IFNL1 (**B**), DENV2 RNA (**C**) by RT-qPCR. hpi, hours post-infection. (**D-F**) MDA5 or RIG-I single knockout cells, as well as DKO HEK293 cells were infected with DENV2 (MOI=1) for 48 h and analyzed by RT-qPCR to detect the mRNA abundance of IFNB1 (**D**), IFNL1 (**E**), DENV2 RNA (**F**). IFNB1 and IFNL1 were normalized against cellular GAPDH mRNA level and expressed as values relative to the mock infection control. DENV2 RNA was expressed as values normalized to the GAPDH mRNA level. The error bars represent standard deviations from two independent experiments. Student’s t test was used for statistical analysis. ns, P>0.05; **, p < 0.01; ***, p < 0.001.

To evaluate the role of MDA5 or RIG-I in innate immune sensing of DENV infection, we generated MDA5 or RIG-I knockout HEK293 cells using the CRISPR-Cas9 technology(32). The loss of MDA5 or RIG-I expression in the knockout cells were verified by the Western blot (Fig. S2A). Next, we transfected these knockout cells with long poly(I:C) or HCV 3’UTR RNA, which are known to be recognized by MDA5 or RIG-I, respectively(15, 16, 33, 34). As expected, poly(I:C)-induced IFN response was decreased in the MDA5 knockout cells but not in the RIG-I knockout cells, whereas HCV 3’UTR RNA-induced IFN response was impaired only in the RIG-I knockout cells (Fig. S2B-2C). Subsequently, the MDA5 and RIG-I knockout HEK293 cells were infected with DENV2 at a MOI of 1 and the IFN production were determined at 48 hours post-infection by RT-qPCR. The knockout of MDA5 had no apparent effect on interferon production, but the knockout of RIG-I significantly impaired the induction of IFNB1 and IFNL1 (Fig. S2D-2E), despite that viral RNA replication was comparable in the three groups of cells (Fig. S2F). These results suggested RIG-I was the main innate sensor for DENV2 in HEK293 cells.

Although the knockout of RIG-I dramatically impaired the IFN response induced by DENV2 infection, considerable IFN signaling was still activated in the RIG-I knockout cells (Fig. S2D-2E). We speculated that MDA5 may also contribute to DENV2 infection-triggered IFN signaling albeit to much less extent than RIG-I. As the result, the MDA5 single knockout does not lead to a significant decrease in IFN production, and the contribution of MDA5-mediated signaling may become more pronounced in the RIG-I deficient cells. To test our hypothesis, we knocked out MDA5 in the RIG-I knockout HEK293 cells to generate double knockout (DKO) cells (Fig. S2G). Next we transfected poly(I:C) or HCV 3’UTR RNA into the RIG-I knockout or DKO cells. The result showed that poly(I:C)-induced IFN response was significantly decreased in the DKO cells, while residual HCV 3’UTR RNA-induced IFN response likely resulted from incomplete RIG-I knockout remained unchanged in the DKO cells (Fig. S2H-2I). Next, the MDA5 or RIG-I single knockout and DKO cells were infected with DENV2, and the IFN responses were detected at 48 hours post-infection. Interestingly, the IFN signaling was further reduced in the DKO cells compared to the RIG-I single knockout cells (Fig. 1D-1E), whereas the viral replication levels were comparable in all four group of cells (Fig. 1F). These results suggested DENV2 infection-induced IFN response was dependent upon both MDA5 and RIG-I, with RIG-I playing a predominant role.

### RIG-I and MDA5 recognize the different moiety of DENV2 replicative-form dsRNA

To identify the nature of PAMPs recognized by RIG-I and MDA5 during DENV2 infection, we sought to purify the DENV PAMPs from total RNAs in the DENV2-infected cells through biochemical approaches which have been used for research on Picornavirus(35), influenza A virus and Sendai virus(36). First, DENV2-infected cellular RNA was extracted from HEK293, Huh7 and Vero cells, and was transfected into HEK293 cells. All of the DENV2-infected cellular RNA induced strong IFN response, whereas the uninfected cellular RNA elicited no IFN production (Fig. 2A-2B). The RNAs extracted from DENV-infected Huh7 cells seemed to possess the most potent ability to trigger IFN response, therefore, we chose DENV2-infected Huh7 cellular RNA in subsequent purification experiments. Next, we assessed whether DENV2-infected cellular RNA-activated IFN response recapitulates the RLR dependency of DENV2 infection-induced IFN response. We transfected the DENV2-infected Huh7 cellular RNA into the MDA5 or RIG-I single knockout, as well as DKO cells. Consistent with the DENV2 infection, the IFN response was not decreased in the MDA5 single knockout cells, but was greatly reduced in the RIG-I single knockout cells, and was further reduced in the DKO cells (Fig. 2C-2D), despite that transfected DENV2 RNA levels were comparable in the four groups (Fig. 2E). These results suggested DENV2-infected cellular RNA indeed contained DENV2 PAMPs recognized by RIG-I and MDA5.

**Fig. 2.**
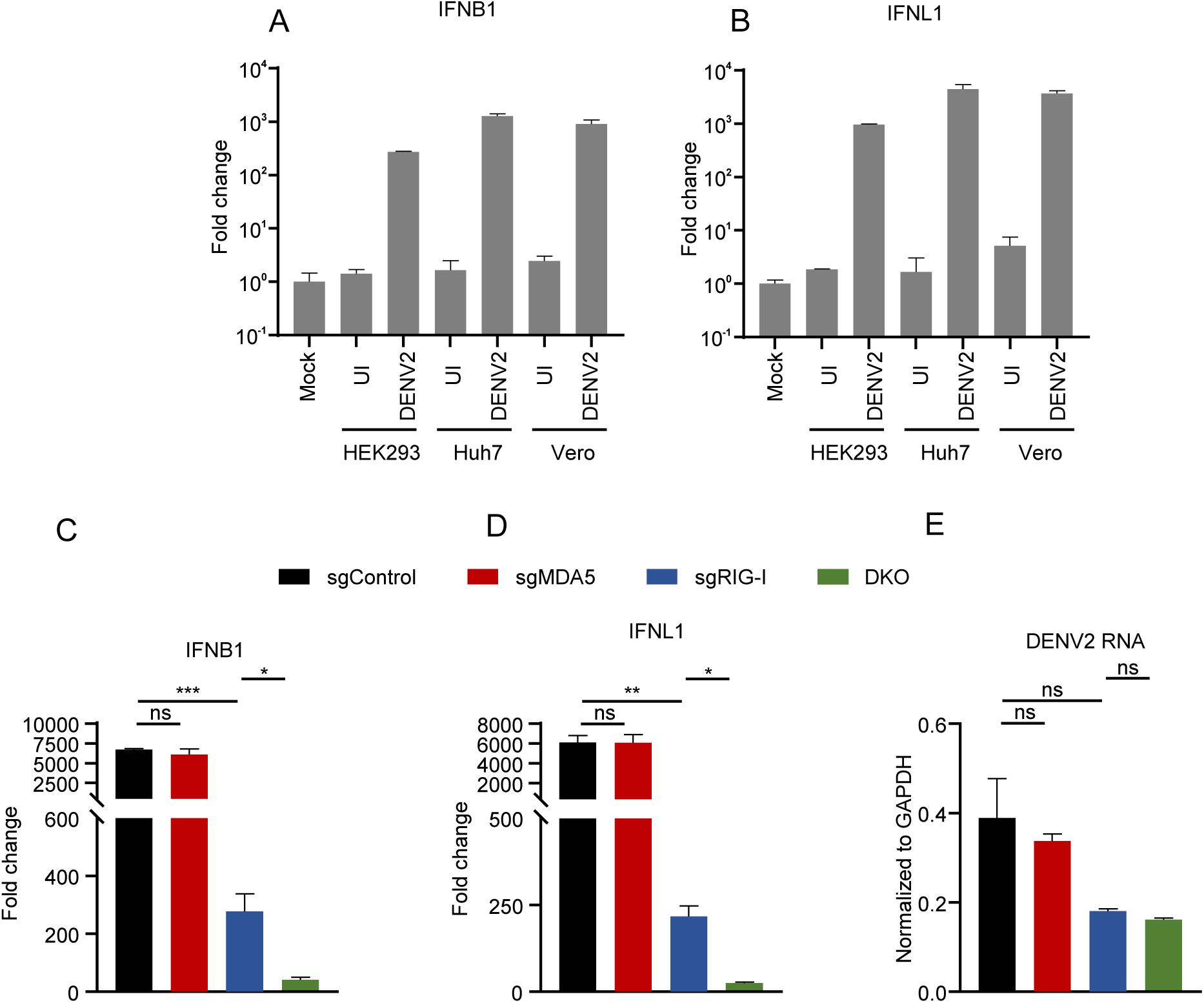
The IFN response stimulated by transfection of DENV2-infected cellular RNA requires both RIG-I and MDA5, with RIG-I playing a predominant role. (**A-B**) HEK293 cells were transfected with the 200 ng of cellular RNA extracted from DENV2 infected or uninfected HEK293, Huh7 and Vero cells for 16 h, and IFNB1 (**A**) and IFNL1 (**B**) mRNA level were determined by RT-qPCR. UI, Uninfected. (**C-E**) RIG-I or MDA5 single knockout cells, as well as DKO HEK293 cells were transfected with 200 ng of DENV2-infected Huh7 cellular RNA for 16 h and analyzed by RT-qPCR to detect the mRNA abundance of IFNB1 (**C**), IFNL1 (**D**), DENV2 RNA (**E**). IFNB1 and IFNL1 were normalized against cellular GAPDH mRNA level and expressed as values relative to the mock transfection control. DENV2 RNA was expressed as values normalized to the GAPDH mRNA level. The error bars represent standard deviations from two independent experiments. Student’s t test was used for statistical analysis for (**C-E**). ns, P>0.05; *, p < 0.05; **, p < 0.01; ***, p < 0.001.

There are four serotypes of DENV (DENV1 to 4)(37). Next, we assessed the IFN activation in the cells infected by other serotypes of DENV. Huh7 cells were infected with DENV1 strain 16007 and DENV4 strain 1036. As shown in Fig. S3A-D, DENV1 and DENV4 infection induced strong interferon responses on day 2 post-infection. We also infected Huh7 cells with DENV3 strain 16562. Consistent with previous studies (38), the replication efficiency of DENV3 was much lower than that of DENV2, and DENV3 infection induced modest IFN responses until day 6 post-infection. Next, the total cellular RNAs from DENV1-, DENV3- and DENV4-infected cells were extracted and subsequently transfected into the MDA5 and RIG-I single knockout, as well as DKO cells. Consistent with the DENV2, DENV1-, DENV3- and DENV4-infected cellular RNA-induced IFN response required both RIG-I and MDA5, with RIG-I playing a primary role (Fig. S4).

To analyze the exact RNA species recognized by MDA5 and RIG-I, we purified PAMPs from the DENV2-infected cellular RNA. First, we separated RNAs according to their sedimentation coefficients via a 10%-40% sucrose density gradient ultracentrifugation (Fig. S5A). The RNA in each fraction was analyzed by agarose gel electrophoresis. The 5S, 18S and 28S ribosomal RNA were distributed in different fractions (Fig. S5B), and DENV2 RNA was primarily distributed in fractions 7-12 (Fig. S5C). To determine which fraction harbored PAMPs, we transfected RNA in each of these 12 fractions into HEK293 cells and analyzed the IFN response. We found that the RNA in fraction number 7, named DENV2-F7, induced the strongest IFN response (Fig. 3A-3B).

**Fig. 3.**
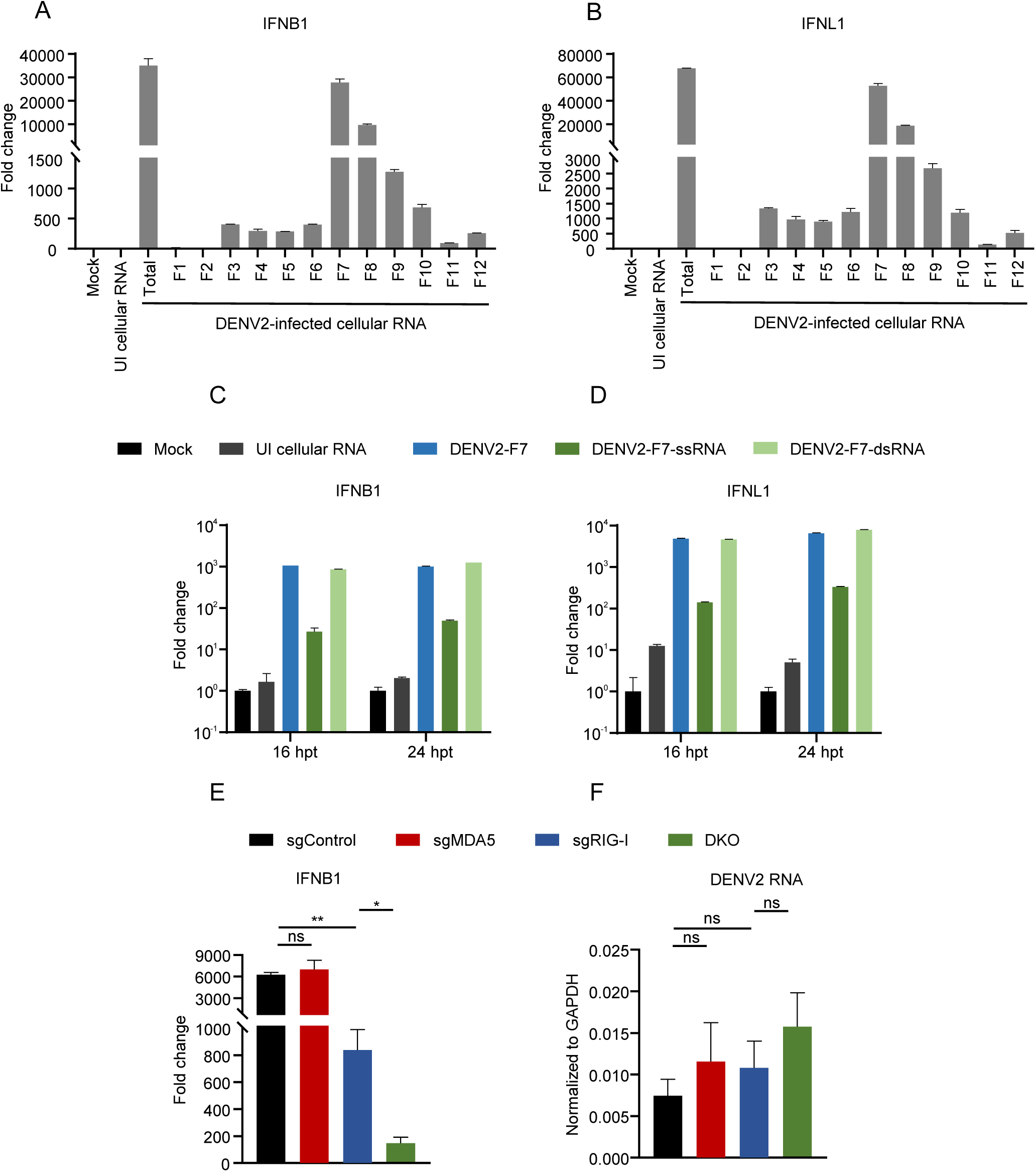
DENV2 PAMPs are enriched through sucrose density gradient centrifugation and LiCl differential precipitation. (**A-B**) HEK293 cells were transfected with equal volume of unfractionated (total) RNA and F1-F12 RNA for 16 h, and IFNB1 (**A**) and IFNL1 (**B**) mRNA levels were determined by RT-qPCR. F1-F12 RNA was stuffed with uninfected Huh7 cellular RNA to make its mass consistent with the unfractionated (total) RNA sample (120 ng). UI, Uninfected. (**C-D**) HEK293 cells were transfected with equal volume of DENV2-F7, DENV2-F7-ssRNA and DENV2-F7-dsRNA. Transfected cells were analyzed at indicated timepoints for IFNB1 (**C**) and IFNL1 (**D**) by RT-qPCR. DENV2-F7 (20 ng), ssRNA and dsRNA were stuffed with 200 ng of uninfected Huh7 cellular RNA to ensure high transfection efficiency. UI, Uninfected. (**E-F**) MDA5 or RIG-I single knockout cells, as well as DKO HEK293 cells were transfected with 0.5 ng of DENV2-F7-dsRNA (stuffed with 200 ng uninfected Huh7 cellular RNA) for 16 h and analyzed by RT-qPCR to detect the mRNA abundance of IFNB1 (**E**) and DENV2 RNA **(F**). IFNB1 and IFNL1 were normalized against cellular GAPDH mRNA level and expressed as values relative to the mock transfection control. DENV2 RNA was expressed as values normalized to the GAPDH mRNA level. The error bars represent standard deviations from two independent experiments. Student’s t test was used for statistical analysis for (**E-F**). ns, P>0.05; *, p < 0.05; **, p < 0.01.

Next, to further purify PAMPs in DENV2-F7, we separated single stranded RNA (ssRNA) and dsRNA by LiCl differential precipitation (Fig. S6A). The quantification of DENV2 RNA result showed that more viral RNA in fraction 7 was found in the dsRNA fraction than the ssRNA fraction (Fig. S6B), whereas majority of viral RNA in fraction 12 was distributed in the ssRNA fraction (Fig. S6C). Then we transfected DENV2-F7 ssRNA and dsRNA into HEK293 cells and analyzed the IFN response. The results showed that dsRNA (DENV2-F7-dsRNA) triggered stronger IFN response than the ssRNA fraction (DENV2-F7-ssRNA) (Fig. 3C-3D). To assess the RLR dependency, we transfected DENV2-F7-dsRNA into MDA5 or RIG-I single knockout, as well as DKO HEK293 cells. Consistent with DENV2-infected cellular RNA, DENV2-F7-dsRNA was capable of eliciting IFN response through both RIG-I- and MDA5-dependent pathways, with RIG-I playing a main role (Fig. 3E-3F), suggesting that purified DENV2-F7-dsRNA likely represents DENV2 PAMPs in the DENV2-infected cells.

Next, we performed a strand-specific RNA sequencing analysis on DENV2-F7-dsRNA. Approximately 42% of reads were mapped to DENV2 genomes, and the DENV RNA was greatly enriched in the preparation of DENV2-F7-dsRNA (Fig. S7), suggesting that the DENV PAMPs are likely derived from viral RNA rather than host RNA. DENV sequencing reads were mapped to their positions on the viral genomes. In DENV2-infected cellular RNA, the RNA reads mapping to the (+) sense strand were significantly greater than those mapping to the (-) sense strand (Fig.4A), consistent with a previous finding that positive RNA viruses tend to produce more (+) sense-strand RNA than (-) sense-strand RNA(39). However, in DENV2-F7-dsRNA, the RNA reads mapping to the both strands were approximately equal across the genome (Fig. 4B). We calculated the ratio of reads mapped to the (+) sense versus those mapped to the (-) sense RNA at each position, expressing this ratio on the log10 scale (Fig. 4C). For DENV2-infected cellular RNA, the average ratio was around 45. However, for DENV2-F7-dsRNA, the ratios were close to 1. The significant differences in read depth mapped to the (+) and (-) sense RNA observed at the ends of genomes might be attributed to a technical bias in sequencing the terminal regions of genes.

**Fig. 4.**
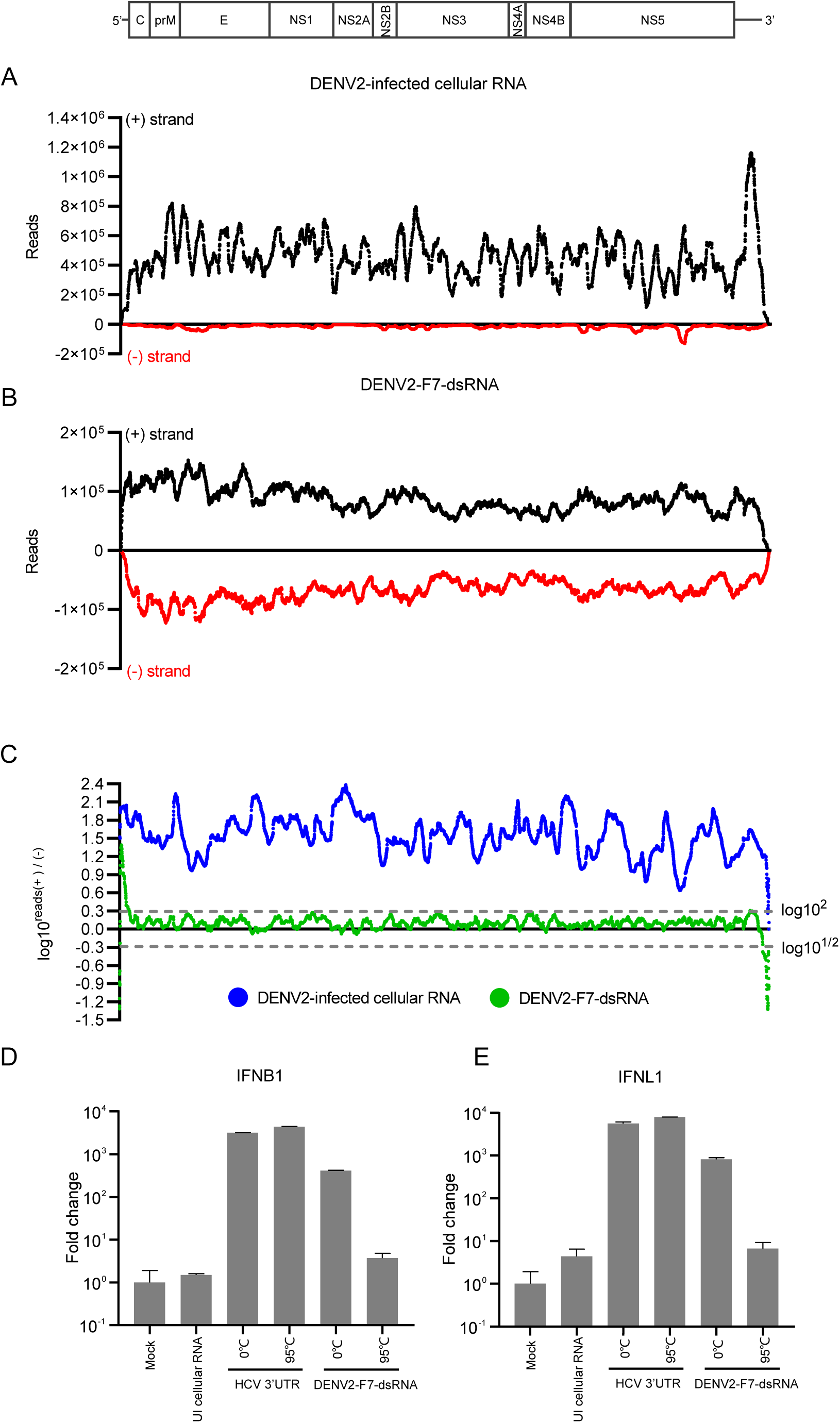
DENV2 PAMPs are the DENV2 RF double-stranded RNA. (**A**) DENV2-infected cellular RNA and (**B**) DENV2-F7-dsRNA samples were subjected to strand-specific RNA-seq analysis. The distribution of read coverage matching DENV2 genomes was represented along the viral genome. The x-axis denoted the nucleotide positions within the DENV2 genome, while the y-axis illustrated the depth of reads aligned to the DENV2 (+) sense RNA in black and the (-) sense RNA in red. (**C**) The ratio was determined by dividing the depth of reads mapped to the (+) sense RNA by the depth of reads mapped to the (-) sense RNA, and the results were represented on a log10 scale. Values between the two gray dotted lines indicated ratios that are within a twofold difference. (**D-E**) 200 ng of HCV 3’UTR and 0.3 ng of DENV2-F7-dsRNA were heated at 95°C for 5 minutes and then transfected into HEK293 cells for 16 h, and IFNB1 (**D**) and IFNL1 **(E**) were analyzed by RT-qPCR. 200 ng of uninfected cellular Huh7 RNA as a carrier were transfected together with heated DENV2-F7-dsRNA. UI, Uninfected. IFNB1 and IFNL1 were normalized against cellular GAPDH mRNA level and expressed as values relative to the mock transfection control. The error bars represent standard deviations from two measurements.

Rapid cooling after high-temperature heating can resolve the duplex of a dsRNA. To further validate the dsRNA feature of DENV2 PAMPs, we heated DENV2-F7-dsRNA at 95℃ for 5 min and rapidly cooled it to 0℃, then transfected them into HEK293 cells. The single-stranded HCV 3’UTR RNA was included as a control in this experiment. As shown in Fig. 4D-E, heat-denaturing treatment abolished the capacity of DENV2-F7-dsRNA to elicit IFN response, but had little effect on the HCV 3’UTR RNA-induced IFN response. Altogether, these results suggested that the DENV PAMPs was likely derived from the viral replicative-form (RF) double-stranded RNA.

The 5’ppp modification of RNA is essential for recognition by RIG-I. Therefore, next we investigated whether DENV2 PAMPs-induced IFN response requires 5’ppp. DENV2-F7-dsRNA was treated with a 5’-polyphosphatase (PPase) to remove the γ- and β-phosphates and then transfected into HEK293 cells.

After treatment of PPase, DENV2-F7-dsRNA-induced IFN production was greatly reduced (Fig. 5A-5B), while the transfected DENV2-F7-dsRNA levels in the cells remained comparable before and after treatment with PPase (Fig. 5C). These results indicated that DENV2 PAMPs triggered-IFN production required 5’ppp modification. We observed that PPase-treated DENV2-F7-dsRNA still retained considerable capacity to induce IFN production (Fig. 5A-5B). It has been shown that recognition of dsRNA by MDA5 is independent on the 5’ppp of dsRNA(40). Therefore, we speculated that PPase-treated DENV2-F7-dsRNA might induce the IFN response exclusively through MDA5. To test our hypothesis, we transfected DENV2-F7-dsRNA with or without PPase treatment into MDA5 or RIG-I single knockout HEK293 cells. Consistent with previous results (Fig. 3E), DENV2-F7-dsRNA-induced IFN response was primarily dependent on RIG-I. However, the PPase treatment converted this IFN induction into a MDA5-dependent manner (Fig. 5D-5F).

**Fig. 5.**
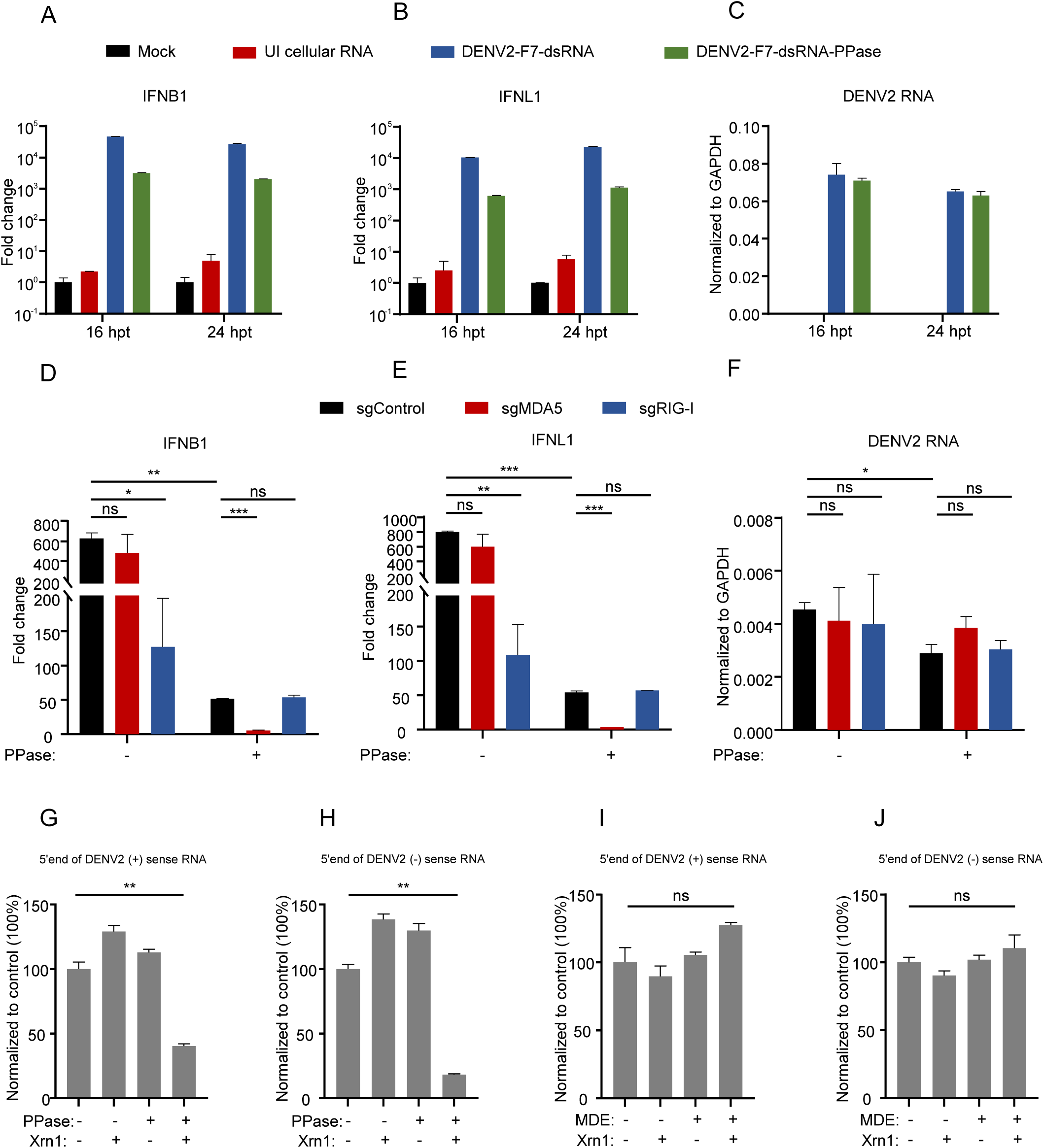
5’ppp modification of DENV2-F7-dsRNA is essential for RIG-I-mediated signaling, but not for MDA5-mediated signaling. (**A-C**) 0.5 ng of PPase-treated or untreated DENV2-F7-dsRNA was stuffed with 200 ng of uninfected Huh7 cellular RNA, and then transfected into HEK293 cells. Transfected cells were analyzed at indicated timepoints for IFNB1 (**A**) and IFNL1 (**B**) and DENV2 RNA **C**) by RT-qPCR. UI, Uninfected. hpt, hours post-transfection. The error bars represent standard deviations from two independent measurements. (**D-F**) RNAs in (**A-C**) were transfected into MDA5 or RIG-I single knockout cells. Transfected cells were analyzed at 16 h for IFNB1 (**D**) and IFNL1 (**E**) and DENV2 RNA (**F**) by RT-qPCR. IFNB1 and IFNL1 were normalized against cellular GAPDH mRNA level and expressed as values relative to the mock transfection control. DENV2 RNA was expressed as values normalized to the GAPDH mRNA level. The error bars represent standard deviations from two independent experiments. (**G-H**) DENV2-F7-dsRNA supplemented with uninfected Huh7 cellular RNA, was subjected to treatment with Xrn1, PPase, or both. DENV2 (+) (**G**) or (-) (**H**) sense RNA were detected by RT-qPCR targeting the 5’end of RNA. **(I-J**) DENV2-F7-dsRNA supplemented with uninfected Huh7 cellular RNA, was subjected to treatment with Xrn1, MDE, or both. DENV2 (+) (**I**) or (-) (**J**) sense RNA were detected by RT-qPCR targeting the 5’end of RNA. The RNA levels were normalized to a control sample. The error bars represent standard deviations from two independent experiments. Student’s t test was used for statistical analysis for (**D-J**). ns, P>0.05; *, p < 0.05; **, p < 0.01; ***, p < 0.001.

Altogether, these results suggested that recognition of DENV PAMPs by RIG-I is dependent on both double-stranded RNA structure and 5’ppp, while recognition of DENV PAMPs by MDA5 only requires double-stranded RNA structure.

The mature DENV genomic (+) sense RNA is capped, a process that relies on the presence of stem loop A (SLA) within the 5’UTR. DENV (-) sense RNA, which lacks SLA, remains uncapped(41). However, it remains uncertain whether the (+) sense RNA within the RF is capped. In West Nile virus (WNV)-infected cells, the cap is shown to be present only on the genomic (+) sense RNA but not on the RF(42). To explore whether the 5’ ends of the DENV RF (+) and (-) sense RNA were modified with 5’ppp, we treated DENV2-F7-dsRNA with PPase to remove the γ- and β-phosphates and then digested the RNA with 5’–3’ exonuclease Xrn1, which digests RNA with a 5’monophosphate (5’p). After treatment with PPase and Xrn1, we assessed the RNA degradation by RT-qPCR using a primer set targeting the 5’ region of the DENV2 (+) stand and (-) strand respectively (Fig. S8A). As controls for the assay, we also included *in vitro* transcribed (IVT) replicon (+) sense ssRNA bearing a 5’ppp and uninfected Huh7 cellular RNAs containing 5’ capped ACTB mRNA. As expected, the PPase treatment rendered the IVT DENV2 replicon ssRNA but not capped ACTB mRNA susceptible to digestion by Xrn1 (Fig. S8B-8C). Importantly, the PPase treatment rendered both (+) and (-) strands of DENV2-F7-dsRNA susceptible to digestion by Xrn1 (Fig. 5G-5H), suggesting that the both strands of DENV2 RF contained a 5’ppp. To further assess whether the 5’end of (+) sense RNA was capped with m7GpppAm. We treated DENV2-F7-dsRNA with mRNA decapping enzyme (MDE) to remove the cap-0 or cap-1 structure, producing a 5’p end, and then digested the RNA with Xrn1 (Fig. S8D). The uninfected Huh7 cellular RNAs containing ACTB mRNA with a 5’ cap and BCYRN1 RNA with a 5’ppp(43) were included as controls for the assay. As expected, the MDE treatment rendered the capped ACTB mRNA but not BCYRN1 RNA susceptible to digestion by Xrn1 (Fig. S8E-8F). Both (+) and (-) strands of DENV2-F7-dsRNA remained stable upon the MDE-Xrn1treatment (Fig. 5I-5J), suggesting that neither strand of DENV2 RF was capped. Taken together, our results demonstrated that (+) and (-) sense RNA of the DENV2 PAMP dsRNA, similar to WNV RF(42), contained a 5’ppp.

### *In vitro* reconstituted DENV RF dsRNA recapitulates the features of DENV PAMPs

Next, we sought to confirm the nature of DENV PAMPs by reconstituting DENV RF *in vitro*. The (+) and (-) sense DENV2 replicon RNAs bearing a 5’ppp were synthesized by *in vitro* transcription, and then annealed at a ratio of 1:1 to generate a dsRNA mimicking DENV RF, which was confirmed by size-exclusion chromatography (Fig. S9). Next, we investigated whether the reconstituted DENV RF dsRNA displays the similar features of DENV2-F7-dsRNA. We transfected the reconstituted DENV RF dsRNA into MDA5 or RIG-I single knockout, as well as DKO HEK293 cells. As expected, the reconstituted DENV RF induced-IFN response required both MDA5 and RIG-I, with RIG-I playing a predominant role (Fig. 6A-6B). Subsequently, the reconstituted DENV RF dsRNA was treated with PPase to generate 5’p, and then transfected into MDA5 or RIG-I single knockout HEK293 cells. Consistent with DENV2-F7-dsRNA, the treatment with PPase reduced the capacity of the reconstituted DENV RF to induce IFN response and converted the IFN production from a RIG-I-dependent to a MDA5-dependent manner (Fig. 6C-6E). Taken together, these results demonstrated that the reconstituted DENV RF dsRNA exhibited similar characteristics to the purified DENV PAMP.

**Fig. 6.**
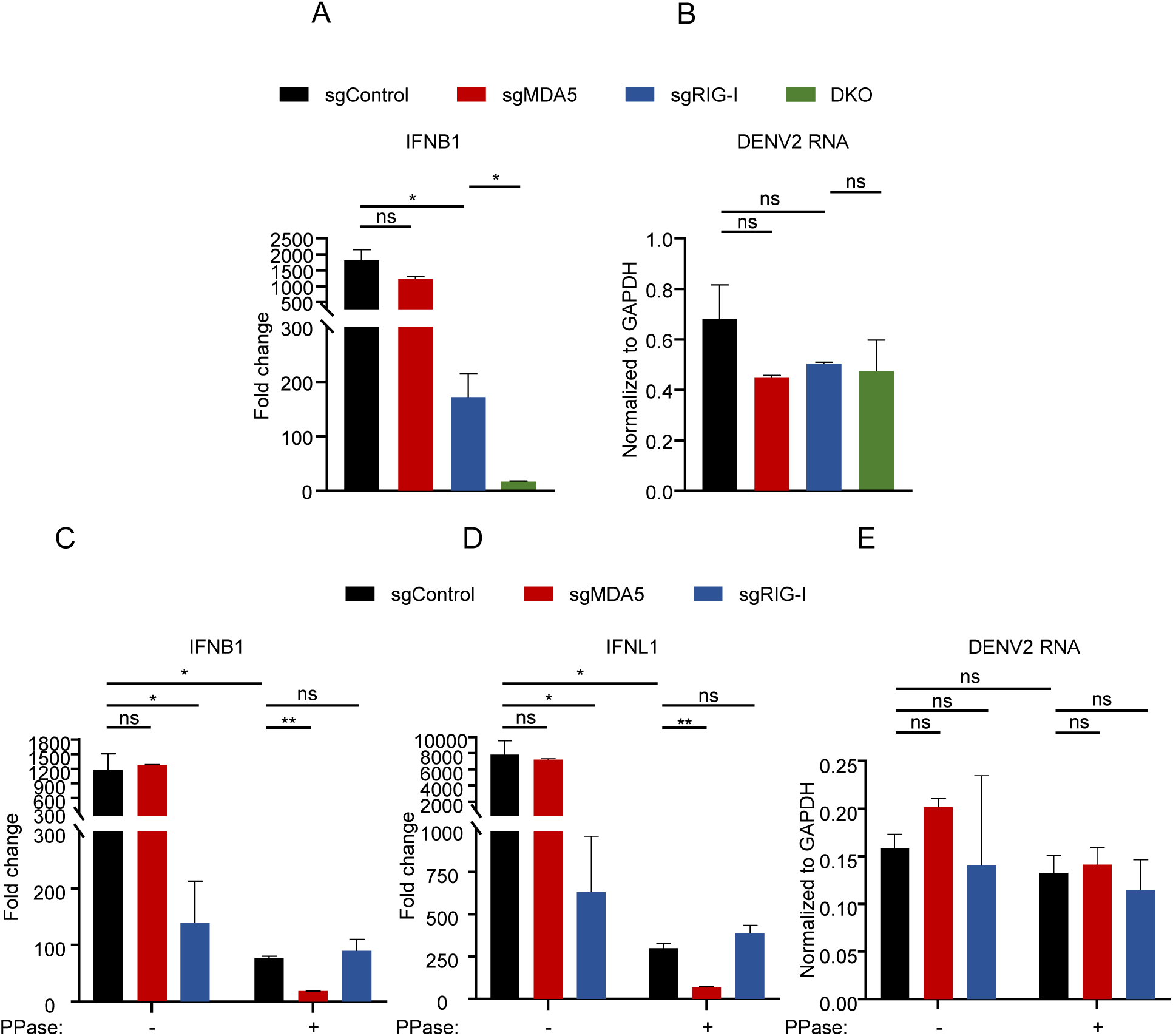
*In vitro* reconstituted DENV2 RF dsRNA exhibits the similar properties to the DENV2 PAMPs. (**A-B**) RIG-I or MDA5 single knockout cells, as well as DKO HEK293 cells were transfected with 50 ng of reconstituted DENV2 RF dsRNA for 16 h and analyzed by RT-qPCR to detect the mRNA abundance of IFNB1 (**A**), DENV2 RNA (**B**). (**C-E**) 50 ng of reconstituted DENV2 RF dsRNAs were treated with PPase, and then transfected into MDA5 or RIG-I single knockout cells. Transfected cells were analyzed at 16 h post-transfection for IFNB1 (**C**) and IFNL1 (**D**) and DENV2 RNA (**E**) by RT-qPCR. IFNB1 and IFNL1 were normalized against cellular GAPDH mRNA level and expressed as values relative to the mock transfection control. DENV2 RNA was expressed as values normalized to the GAPDH mRNA level. The error bars represent standard deviations from two independent experiments. Student’s t test was used for statistical analysis. ns, P>0.05; *, p < 0.05; **, p < 0.01.

## Discussion

Currently, the exact molecular identity of DENV PAMPs recognized by RLRs under physiological conditions has not yet been fully explored. Additionally, it remains controversial which PRR exactly mediates IFN production during DENV infection, in particular, the importance of RIG-I versus MDA5 in sensing DENV PAMP remains a debatable subject. In this study, we uncovered a cooperative contribution of RIG-I and MDA5 in innate immune sensing DENV infection, with RIG-I playing a predominant role. Next, we purified PAMP RNA from DENV-infected cells using sucrose density gradient centrifugation and LiCl differential precipitation. Using strand-specific RNA-seq, heat denaturing, 5’ end RNA modification-specific enzymatic digestions as well as *in vitro* reconstitution, we demonstrated that the DENV PAMP is likely a DENV RF dsRNA bearing 5’ppp modifications at the both strands.

We found that single RIG-I knockout led to a significant decrease in IFN production during DENV infection, whereas single MDA5 knockout had no effect. However, the function of MDA5 became more pronounced in the RIG-I deficient cells (Fig. 1D-1F and Fig. S4). Maxime and colleagues(22) showed that RIG-I is the main RLR sensor of DENV2 and DENV4 infections in HEK293 cells, whereas diminished expression of MDA5 had little or no effect on IFN production. In HEK293 cells, the contribution of MDA5 to the initiation of the IFN response during DENV infection is indeed quite modest and could be overlooked. Joris K and colleagues(20, 21) reported that silencing RIG-I, MDA5 alone or together strongly abrogated the IFNs and cytokines expression upon DENV2 infection in human dendritic cells. The relative contributions of RIG-I versus MDA5 to the IFN response in the context of these RNA virus infections may vary across different cell types, likely due to the disparities in the relative strength of the RIG-I and MDA5 signaling pathways as well as the availability of corresponding PAMPs in these cells.

Maxime and colleagues(22) proposed that RIG-I detected 5’ppp nascent (+) sense RNA of the DENV RI during viral infection. It is reported that DENV RI sediments between 20 and 28S, and insoluble in 2 M LiCl(39). According to these characteristics, DENV RI is likely to be distributed in F7 to F9 in our sucrose density gradient centrifugation, and enriched in a ssRNA preparation by LiCl differential precipitation.

However, compared to F7 and F7-dsRNA samples, F8, F9 and F7-ssRNA samples induce much less IFN response (Fig. 3). In addition, Nicolas and colleagues(44) reported that RIG-I detected RNA polymerase III (Pol3)-transcribed Y RNA and vault RNA (vtRNA) during DENV4 infection. However, our results showed that viral RNAs were significantly enriched in the DENV2-F7-dsRNA preparation, strongly supporting that DENV PAMPs are likely viral RNAs instead of host cellular RNAs. Taken together, although our results do not completely rule out a minor role of DENV RI, Y RNA and vtRNA in activating IFN response during DENV infection, the major DENV PAMPs are derived from DENV RF dsRNA.

Our work suggested that RIG-I and MDA5 may act simultaneously to sense DENV RF dsRNA despite that these two PRRs tend to recognize a dsRNA with the different molecular feature. The RF bearing 5’ppp may exist during replication of other positive-stranded RNA viruses, for example HCV(43) and WNV(42). Hence, the cooperative detection of PAMP by RIG-I and MDA5 is likely a more general phenomenon during RNA virus infections. Feng and colleagues (35) reported that MDA5 but not RIG-I detected dsRNA RF during Picornavirus infection. This may be due to the fact that the (+) sense and (-) sense strands of Picornaviral RF dsRNA are covalently linked with VPg, a small peptide of 20–24 amino acids that serves as a primer for RNA synthesis(45), thereby preventing them being recognized by RIG-I. Consequently, the Picornavirus RF lacking 5’ppp exclusively activates MDA5 but not RIG-I.

RLRs recognize dsRNA-like structures to activate the IFN response during RNA virus infection. However, the exact nature of these dsRNA-like structures generated during RNA virus infection remains an important subject to explore for many RNA viruses. A number of studies explored RLRs-associated RNAs by precipitating overexpressed or endogenous RLRs, followed by next-generation sequencing(22, 36, 44, 46–49). However, this approach is critically dependent on the efficiency and specificity of the RNA precipitation process. In addition, RLRs-bound RNAs may have undergone irreversible conformational changes upon the RLR binding that lead to loss of the immunostimulatory capability when these RNAs are immunoprecipitated and re-introduced into a cell. In our study, we purified PAMPs from the virus-infected cells by employing sucrose density gradient centrifugation and LiCl differential precipitation. This approach results in a high recovery rate of PAMPs and ensures the stability of PAMPs throughout the purification procedure. Therefore, our method can also be applied to study the identity of PAMPs of other RNA viruses in the context of their infection.

It is well known that DENV (+) sense RNA is capped, but it remains to be elucidated at which point during its synthesis this capping reaction occurs. Valerie J and colleagues proposed two models: co-transcriptional model and post-transcriptional model(4). In the co-transcriptional model, as the nascent (+) sense RNA is transcribed from the DENV RF, the RNA is capped. Following the synthesis of a short stretch of the (+) sense RNA, the 5’ end of this newly synthesized RNA is capped by NS3 and NS5 proteins. Subsequently, the 5’capped (+) sense RNA is extended until the entire strand is fully replicated. Ultimately, a dsRNA containing a capped (+) sense and a free capped (+) sense RNA are produced. In post-transcriptional model, capping takes place on the pre-existing full-length (+) sense but not nascent (+) sense RNA. After the double-stranded RF is unwound by the NS3 helicase, the (+) sense RNA is capped. In the end, a new dsRNA and a free capped (+) sense RNA are synthesized. The primary distinction between the two models is whether the (+) sense RNA within the RF is capped. In our study, we found that both (+) and (-) sense RNA of RF were modified with 5’ppp, and this result provided an important piece of evidence to support the post-transcriptional model.

DENV infection has the potential to induce viral hemorrhagic fever, which is characterized by increased vascular permeability, hemorrhage and shock. Anon and colleagues proposed(50) that severe dengue is mediated by multiple immune effectors. As of now, there are no specific antiviral agents for dengue, treatment relies on fluid replacement and the administration of analgesics to alleviate symptoms. Our study identified RIG-I and MDA5 cooperatively detected the DENV RF during viral infection, providing a potential therapeutic target for the treatment DENV hemorrhagic fever.

## Materials and methods

### Cell culture and virus preparation

HEK293, Huh7, Vero cells were purchased from the National Collection of Authenticated Cell Cultures (Shanghai, China). These cells and their derivatives were maintained in complete Dulbecco’s modified Eagle’s medium (DMEM) (Invitrogen, Carlsbad, CA, USA) supplemented with 10% fetal bovine serum, 10 mM HEPES, 2 mM L-glutamine, 100 U of penicillin/ml, and 100 mg of streptomycin/ml. All cells were cultured in humidified air containing 5% CO2 at 37℃. DENV1/2/3/4 viruses were obtained from Dr. Xia Jin (Institut Pasteur of Shanghai, Chinese Academy of Sciences). All DENV viruses were propagated in Vero cells and titrated by immunofluorescence assay.

### Reagents

Long poly(I:C) was purchased from Invivogen (San Diego, CA). Preparation of HCV 3’UTR RNA was as previously described(17). 5’ polyphosphatase was purchased from Epicentre (RP8092). The mRNA decapping enzyme was purchased from New England Biolabs (M0608S). Xrn1 was purchased from New England Biolabs (M0338S).

### Generation of RIG-I- and MDA5-knockout HEK293 cells by lenti-CRISPR-Cas9

The sequence of sgRNA targeting RIG-I was: 5’-GGCCATGTAGCTCAGGATGT-3’; sgRNA targeting MDA5 was 5’-CGAATTCCCGAGTCCAACCA-3’; sgRNA targeting control was 5’- AACCGCATCGAGCTGA-3’. HEK293T cells were seeded into six-well plates 1 day before transfection and then cotransfected with 1.5 μg of psPAX2 plasmid, 1 μg of pMD2.G plasmid, and 2 μg of lenti-Cas9-sgRNAs using polyethylenimine (Polysciences). Culture supernatants containing lentiviruses were harvested at 48 hours post-transfection, passed through a 0.45-μm-pore-size filter, and used to infect HEK293 cells. Transduced cells were selected in medium containing 1 μg/mL of puromycin for 2 days. To generate DKO cells, RIG-I knockout HEK293 cells were infected with lentiviruses expressing sgMDA5 harboring blasticidin resistance gene. As control, sgControl, sgRIG-I and sgMDA5 cells were infected with lentiviruses expressing sgControl harboring blasticidin resistance gene. Transduced cells were selected in medium containing 15 μg/mL of blasticidin for 2 days. The knockout efficiency was verified by Western blot.

### RNA isolation and quantitative RT-PCR (RT-qPCR)

The protocol was as described previously(51). In brief, total cellular RNA was isolated using TRNzol reagent (Tiangen, Beijing, China). RNA heated at 95°C for 5 minutes and immediately placed on ice to cool down to 0°C, followed by RT-qPCR. RT-qPCR was carried out utilizing SYBR green real-time PCR master mix (Toyobo). The sequences of the qPCR primers for GAPDH and IFNL1 were described previously(52). Other qPCR primers were as follows: IFNB1-F, 5’- GTCACTGTGCCTGGACCATAG-3’; IFNB1-R, 5’-GTTTCGGAGGTAACCTGTAAGTC-3’.

DENV2 RNA-F, 5’-AGTAGTTAGTCTACGTGGAC-3’; DENV2 RNA-R, 5’- GTTGACACGCGGTTTCTCTC-3’. DENV1 RNA-F, 5’-AGTTGTTAGTCTACGTGGAC-3’; DENV1 RNA-R, 5’-GTTGACACGCGGTTTCTCGC-3’. DENV3 RNA-F, 5’-AGTTGTTAGTCTACGTGGAC-3’; DENV3 RNA-R, 5’- GTTGACACACGGTTTCTCAC-3’. DENV4 RNA-F, 5’-AGTTGTTAGTCTGTGTGGAC-3’; DENV4 RNA-R, 5’-GTTGATACGCGGTTTCTCTC-3’. qPCR primers for GAPDH used in Fig. S1 (target both human and green monkey) were as follows: GAPDH-F, 5’-GCTGAGTACGTCGTGGAGTC-3’; GAPDH-R, 5’-TGATGACCCTTTTGGCTCCC-3’.

The qPCR primers used in 5’ end RNA modification-specific enzymatic digestions (Fig. S8 and Fig.5) were as follows: ACTB-F, 5’-GCCTCGCCTTTGCCGA-3’; ACTB-R, 5’-AGGAATCCTTCTGACCCATGC-3’. BCYRN1-F, 5’-GAGGATAGCTTGAGCCCAGG-3’; BCYRN1-R, 5’-GCTTTGAGGGAAGTTACGCTTAT-3’. The qPCR primers for 5’end of DENV2 (+) sense RNA were same as primers for DENV2 RNA as mentioned above. The qPCR primers for 5’end of DENV2 (-) sense RNA: F, 5’-TTGAGTAAACTGTGCAGCCTGTAGCTC-3’; R, 5’-GAGACAGCAGGATCTCTGGTCTTTC-3’.

The qPCR primers used for detecting DENV2 RNA level following sucrose density gradient centrifugation (Fig. S5A) and LiCl differential precipitation (Fig. S6B-6C) were designed to target DENV2 NS3-coding region as follow: DENV2 NS3-F, 5’-AGCATCGAAGACAACCCAGA-3’; DENV2 NS3-R, 5’-CAACTCTAGTGGGGGCCAAG-3’. For the absolute quantification assay (Fig. S7), DENV2 RNA levels in DENV2-infected cellular RNA, DENV2-F7 and DENV2-F7-dsRNA samples were determined relative to a standard curve comprised of serial dilutions of DENV2 replicon plasmid. The qPCR primers were designed to target DENV2 NS3-coding region as mentioned above.

### RNA transfection

One day before transfection, 1×10^5^ HEK293 cells were seeded onto 48-well plates and RNA was transfected using X-TremeGENE HP DNA Transfection Reagent (Roche, Basel, Switzerland) following the manufacturer’s instructions.

### Western blotting

The protocol was as described previously(52). The antibody recognizing MDA5, RIG-I, were obtained from Cell Signaling Technology (Danvers, MA, USA). Antibody against ACTB was obtained from Abmart (Shanghai, China).

### Sucrose density gradient centrifugation

Huh7 cells were infected with DENV2 at MOI of 1. The cells were collected at 48 hpi. Total cellular RNA was extracted using TRNzol reagent. 800 μg (200 μl) DENV2-infected Huh7 cellular RNA were separated by centrifugation in 10%–40% (w/w) sucrose gradient (buffer 20 mM Na-acetate, 100 mM NaCl, 1 mM EDTA, 10U/ml RNasin) at 140,000 ×g in an SW60 Ti rotor (Beckman) at 4°C for 16 h. 12 fractions were collected from top to bottom with 330 μl in each fraction. RNA of 12 fractions were precipitated with isopropanol and then diluted in 200 μl DEPC H2O.

### LiCl differential precipitation

RNA was added with LiCl to a final concentration of 2 M, incubated at 4°C for 16 h. The ssRNA fraction was precipitated by centrifugation at 16,000 ×g for 30 min at 4°C. The supernatant was subjected to ethanol precipitate in the presence of 0.8M LiCl to precipitate the dsRNA fraction. RNA pellets were washed with 75% ethanol, air-dried, and dissolved in DEPC H2O.

### Strand-specific RNA-seq and data analysis

The cDNA library construction and sequencing were conducted by NeoBio, Shanghai, China. DENV2-infected cellular RNA and DENV2-F7-dsRNA were used as the starting material. To analyze all RNA species present, the poly(A) RNA isolation step was omitted. The Ribo-Zero rRNA Removal Kit (Epicentre, USA) was used for DENV2-infected cellular RNA to remove rRNA. The DENV2-F7-dsRNA had a low rRNA content, therefore, the step of removing rRNA was omitted. The Truseq Stranded mRNA sample preparation kit (Illumina,USA) was used for the RNA fragmentation, the first-strand cDNA synthesis, and the second-strand cDNA synthesis. The double strands cDNA was then purified for end repair, dA tailing, adaptors ligation and DNA fragments enrichment. Quantify the libraries using Qubit (invitrogen,CA,USA) according to the Qubit user Guide. The libraries were then pooled and sequenced on NovaSeq 6000 (Illumina). The raw data was handled by fastp (version 0.23.2) and data quality was checked by FastQC (version 0.12.1). Clean reads were aligned to the genome files build with the human genome (hg38) plus DENV2 strain 16681 sequence, and the alignment was performed by bwa (Version 0.7.17).

Samtools (version 1.15.1) was used to extract reads that match the forward and reverse chains, and saved as two new bam files respectively. Finally, featureCounts (v2.0.3) was used to perform the quantification of genes using the above bam files.

### 5’-end RNA modification-specific enzymatic digestion assays

For samples with lower RNA quantity, such as (IVT) DENV2 replicon (+) sense RNA and DENV-F7-dsRNA were supplemented with 10 μg uninfected Huh7 cellular RNA when treated with PPase or MDE. To convert 5’ppp to 5’p, 0.5 U/μl of PPase in 40 μl of 1× reaction buffer (provided by the kit) for 30 min at 37°C followed by purification with TRNzol reagent. To convert 5’cap to 5’p, 5 U/μl of MDE in 20 μl of 1× reaction buffer (provided by the kit) for 60 min at 37°C followed by purification with TRNzol reagent. PPase or MDE treated DENV-F7-dsRNA was denatured by heat (95 °C for 5 min) to generate ssRNA for Xrn1 degradation. For 5’p RNA degradation, RNA was treated with 0.1 U/μl of Xrn1 in 10 μl of 1× NEB buffer 3 for 60 min at 37 °C followed by purification with TRNzol reagent. The RNA degradation efficiency was determined by RT-qPCR.

### *In vitro* reconstitution of DENV2 RF dsRNA

PCR products were used as DNA templates for *in vitro* transcription. The DNA templates were amplified from DENV2 replicon plasmid(53). To amplify DNA templates for DENV2 replicon (+) sense RNA using the following primers, F, 5’-GCCGCAAATTTAATACGACTCACTATAGAGT-3’, and R, 5’-GGTGCTGTTGAATCAACAGGTTCT-3’. To amplify DNA templates for DENV2 replicon (-) sense RNA using the following primers, F, 5’-AGTTGTTAGTCTACGTGGACCGACAAAGACAGATTCT-3’, and R, 5’-GCCGCAAATTTAATACGACTCACTATAGAGAACCTGTTGATTCAACAGCACCATTCCATTTTCT-3’. *In vitro* transcription of DENV2 replicon (+) and (-) sense RNA were performed using a MEGAscript T7 transcription kit (Invitrogen, Carlsbad, CA, USA) following the manufacturer’s protocol. To form a double-stranded substrate, the DENV2 replicon (+) and (-) sense RNA were mixed in equimolar concentrations in 20 mM Tris buffer (pH 8.0) containing 1 mM EDTA and 0.1 M NaCl. The mixture was heated at 95°C for 5 min and then cooled to 30°C for 40 min. The annealed duplex was extracted by TRNzol reagent.

### Size-exclusion chromatography (SEC)

The SEC was performed with a Bio SEC-5 2000 Å (i.d. 7.8 mm; length 300 mm) column (Agilent Technologies, Foster City, CA), with 100 mM ammonium acetate (pH 7.0) as the mobile phase. Isocratic separations were performed at 0.5 ml/min with RNA elution monitored by UV absorbance on the Agilent 1200 HPLC system.

### Statistical analysis

GraphPad Prism 8 software (GraphPad Software Inc.) was used for statistical analysis. Statistical analyses were performed using unpaired, two-tailed Student’s t test. P values <0.05 was considered statistically significant.

## Acknowledgements

We thank Dr. Xia Jin (Institut Pasteur of Shanghai, Chinese Academy of Sciences) for providing DENV1, DENV2, DENV3 and DENV4. We thank Dr. Gong Cheng (Tsinghua University, China) for providing the DENV2 replicon plasmid. We thank Yujiao Qin (Shanghai Institute of Immunity and Infection, Chinese Academy of Sciences) for providing technical support on size-exclusion chromatography. This work was supported by the grants from Key Research and Development Program, Ministry of Science and Technology of China (2021YFC2300204, 2022YFC2303300), National Natural Science Foundation of China (32070944, 32261143735), Program of Shanghai Academic/Technology Research Leader (22XD1403800) to J.Z.

## Author Contributions

S.Y. designed and conducted investigation, analyzed the data, and prepared the manuscript. Y.L., Y.C. and B.L. assisted the investigation. J.Z. conceived the idea, designed investigation, supervised the study and edited the manuscript.

## Data Availability

All data are available upon request.

## Supplementary materials

**Fig S1.**
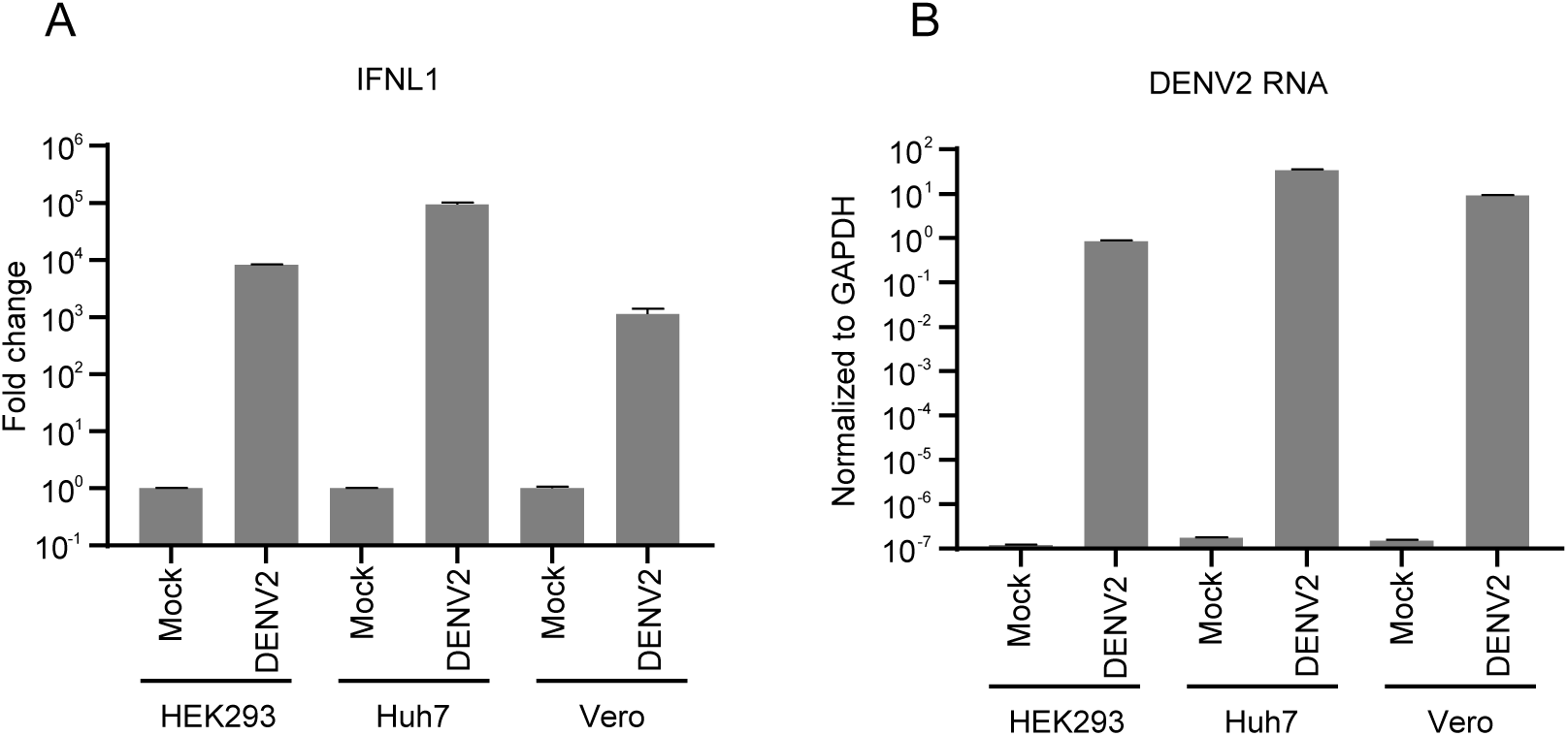
DENV2 infection induces IFN response in HEK293, Huh7 and Vero cells. (**A-B**) HEK293, Huh7 and Vero cells were infected with DENV2 (MOI = 1) for 48 hours and analyzed by RT-qPCR to detect the mRNA abundance of IFNL1 (**A**) and DENV2 RNA (**B**). IFNL1 were normalized against cellular GAPDH mRNA level and expressed as values relative to the mock infection control. DENV2 RNA was expressed as values normalized to the GAPDH mRNA level. The error bars represent standard deviations from two measurements.

**Fig S2.**
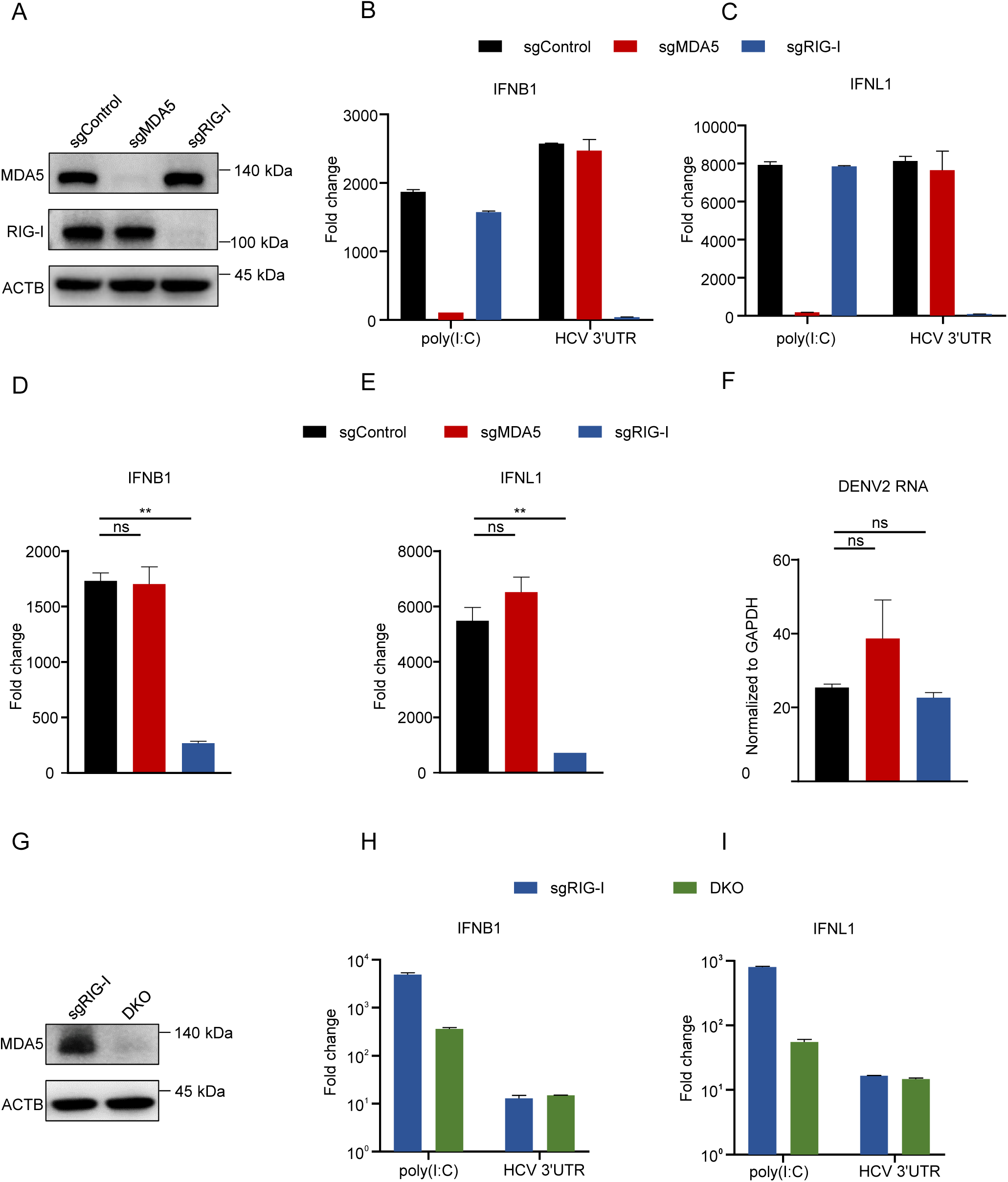
DENV2 infection-induced interferon response is mainly dependent on RIG-I. (**A**) Immunoblotting assay of the MDA5 and RIG-I protein expression in MDA5 or RIG-I single knockout HEK293 cells that had been treated with 10,000 IU/ml IFN-a for 48 h. (**B-C**) The knockout cells were transfected with 200 ng of HCV 3’UTR RNA or 200 ng of long poly(I:C) for 16 h, and IFNB1 (**B**) and IFNL1 (**C**) mRNA level were determined by RT-qPCR. The error bars represent standard deviations from two measurements. (**D-F**) The MDA5 or RIG-I single knockout cells were infected with DENV2 (MOI=1) for 48 h and analyzed by RT-qPCR to detect the mRNA abundance of IFNB1 (**D**), IFNL1 (**E**), DENV2 RNA (**F**). The error bars represent standard deviations from two independent experiments. Student’s t test was used for statistical analysis. ns, P>0.05; **, p < 0.01. (**G**) Immunoblotting assay of the MDA5 expression in the RIG-I single knockout and DKO HEK293 cells that had been treated with 10,000 IU/ml IFN-a for 48 h. (**H-I**) Knockout cells were transfected with the 200 ng HCV 3’UTR RNA or 200 ng long poly(I:C) for 16 h, and IFNB1 (**H**) and IFNL1 (**I**) mRNA level were determined by RT-qPCR. The error bars represent standard deviations from two measurements. IFNB1 and IFNL1 were normalized against cellular GAPDH mRNA level and expressed as values relative to the mock control. DENV2 RNA was expressed as values normalized to the GAPDH mRNA level.

**Fig S3.**
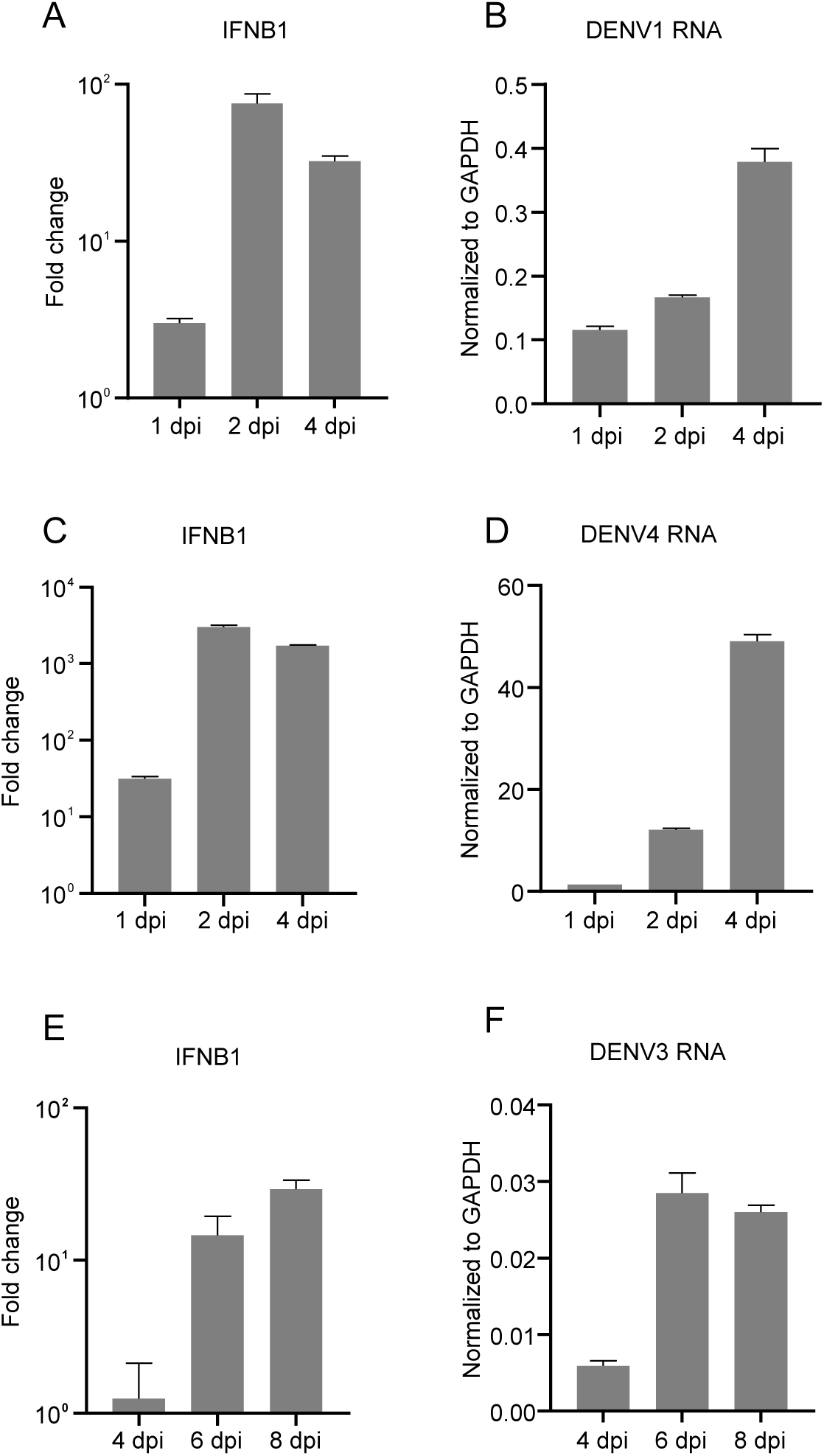
DENV1, DENV3 and DENV4 infection induce IFN response in Huh7 cells. (**A-B**) Huh7 cells were infected with DENV1 (MOI=1). Infected cells were analyzed at indicated time points for IFNB1 (**A**) and DENV1 RNA (**B**) by RT-qPCR. (**C-D**) Huh7 cells were infected with DENV4 (MOI=1). Infected cells were analyzed at indicated time points for IFNB1 (**C**) and DENV4 RNA (**D**) by RT-qPCR. (**E-F**) Huh7 cells were infected with DENV3 (MOI=0.0005). Infected cells were analyzed at indicated time points for IFNB1 (**E**) and DENV3 RNA (**F**) by RT-qPCR. dpi, days post-infection. IFNB1 was normalized against cellular GAPDH mRNA level and expressed as values relative to the mock infection control. DENV1, DENV3 and DENV4 RNAs were expressed as values normalized to the GAPDH mRNA level. The error bars represent standard deviations from two measurements.

**Fig S4.**
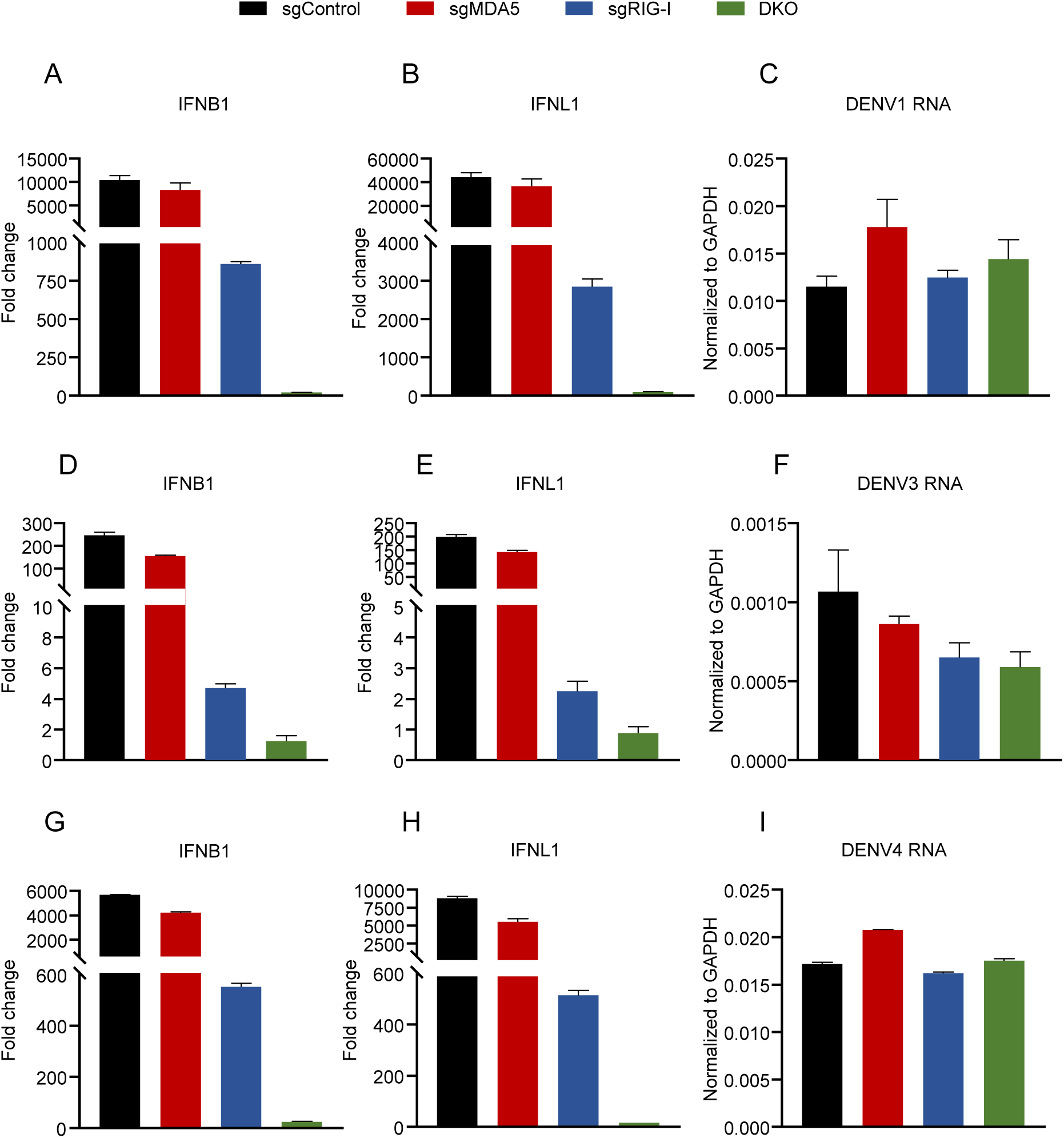
Cellular RNA extracted from DENV1-, DENV3- and DENV4-infected cells induces interferon response which is dependent upon both RIG-I and MDA5, with RIG-I playing a predominant role. (**A-C**) MDA5 or RIG-I single knockout cells, as well as double knockout (DKO) HEK293 cells were transfected with 1μg of DENV1-infected Huh7 cellular RNA (2 dpi) for 16 h, and analyzed by RT-qPCR to detect the mRNA abundance of IFNB1 (**A**), IFNL1 (**B**), DENV1 RNA (**C**). (**D-F**) MDA5 or RIG-I single knockout cells, as well as double knockout (DKO) HEK293 cells were transfected with 1μg of extracted DENV3-infected Huh7 cellular RNA (6 dpi) for 16 h, and analyzed by RT-qPCR to detect the mRNA abundance of IFNB1 (**D**), IFNL1 (**E**), DENV3 RNA (**F**). (**G-I**) The MDA5 or RIG-I single knockout cells, as well as DKO HEK293 cells were transfected with 200 ng of DENV4-infected Huh7 cellular RNA (2 dpi) for 16 h and analyzed by RT-qPCR to detect the mRNA abundance of IFNB1 (**G**), IFNL1 (**H**), DENV4 RNA (**I**). IFNB1 and IFNL1 were normalized against cellular GAPDH mRNA level and expressed as values relative to the mock transfection control. DENV1, DENV3 and DENV4 RNA were expressed as values normalized to the GAPDH mRNA level. The error bars represent standard deviations from two measurements.

**Fig S5.**
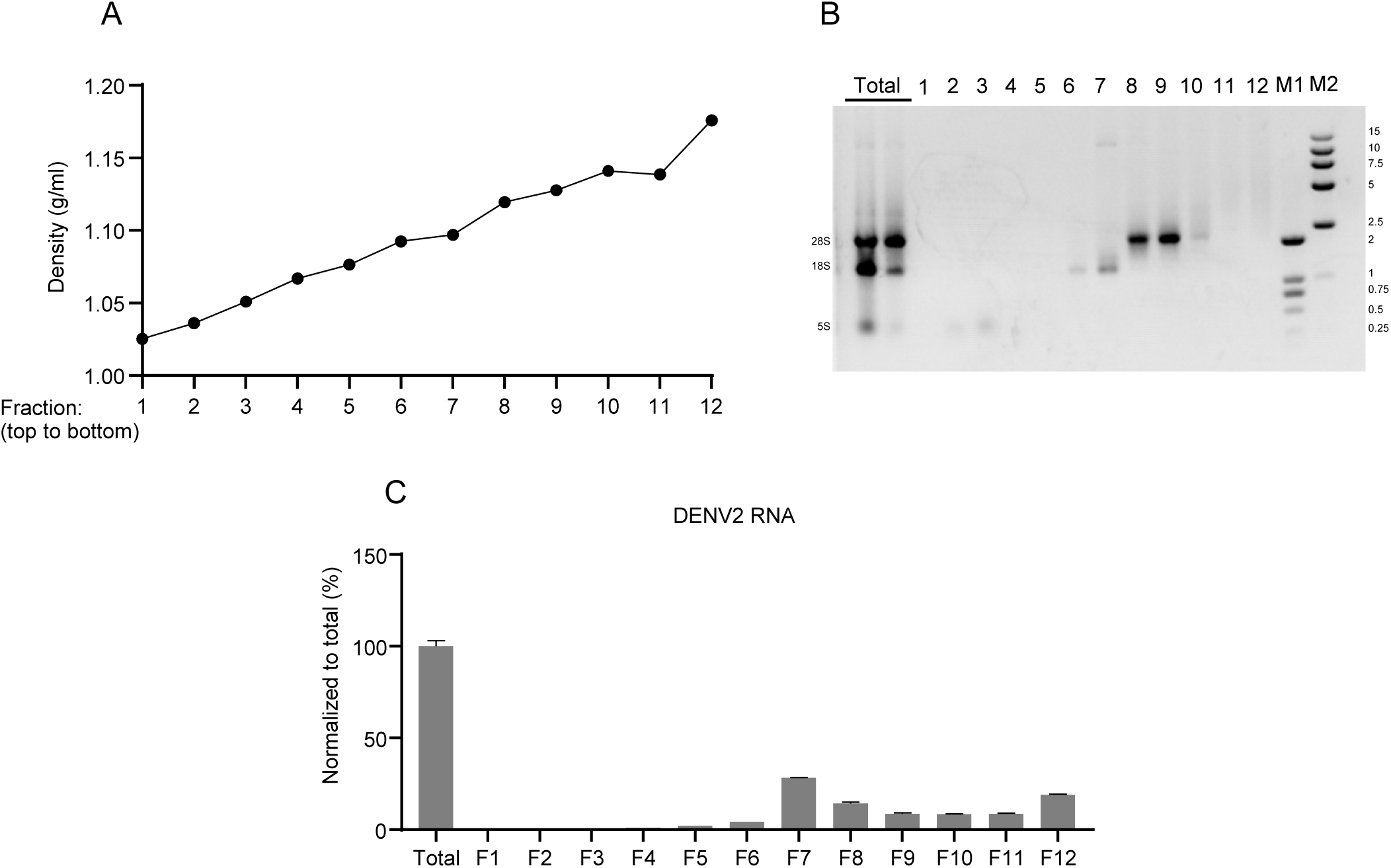
Purification of DENV2 PAMPs from DENV2-infected cells through sucrose density gradient ultracentrifugation. (**A-C**) DENV2-infected cellular RNA was fractionated through sucrose density gradient ultracentrifugation. Fractions (1–12) were collected from the top to bottom of the gradient. (**A**) Densities of 12 fractions were determined. (**B**) The equal volume of unfractionated (total) RNA (line1) and each of 12 fractions (F1-F12) RNA were analyzed by 0.75% of native agarose gel electrophoresis. Total RNA (line 2) was diluted 2.5-fold relative to the total RNA (line 1). M1 and M2 are DNA markers with indicated size in kbp. (**C**) The DENV2 RNA levels in F1-F12 were analyzed by RT-qPCR and normalized to unfractionated (total) sample. The error bars represent standard deviations from two experiments.

**Fig S6.**
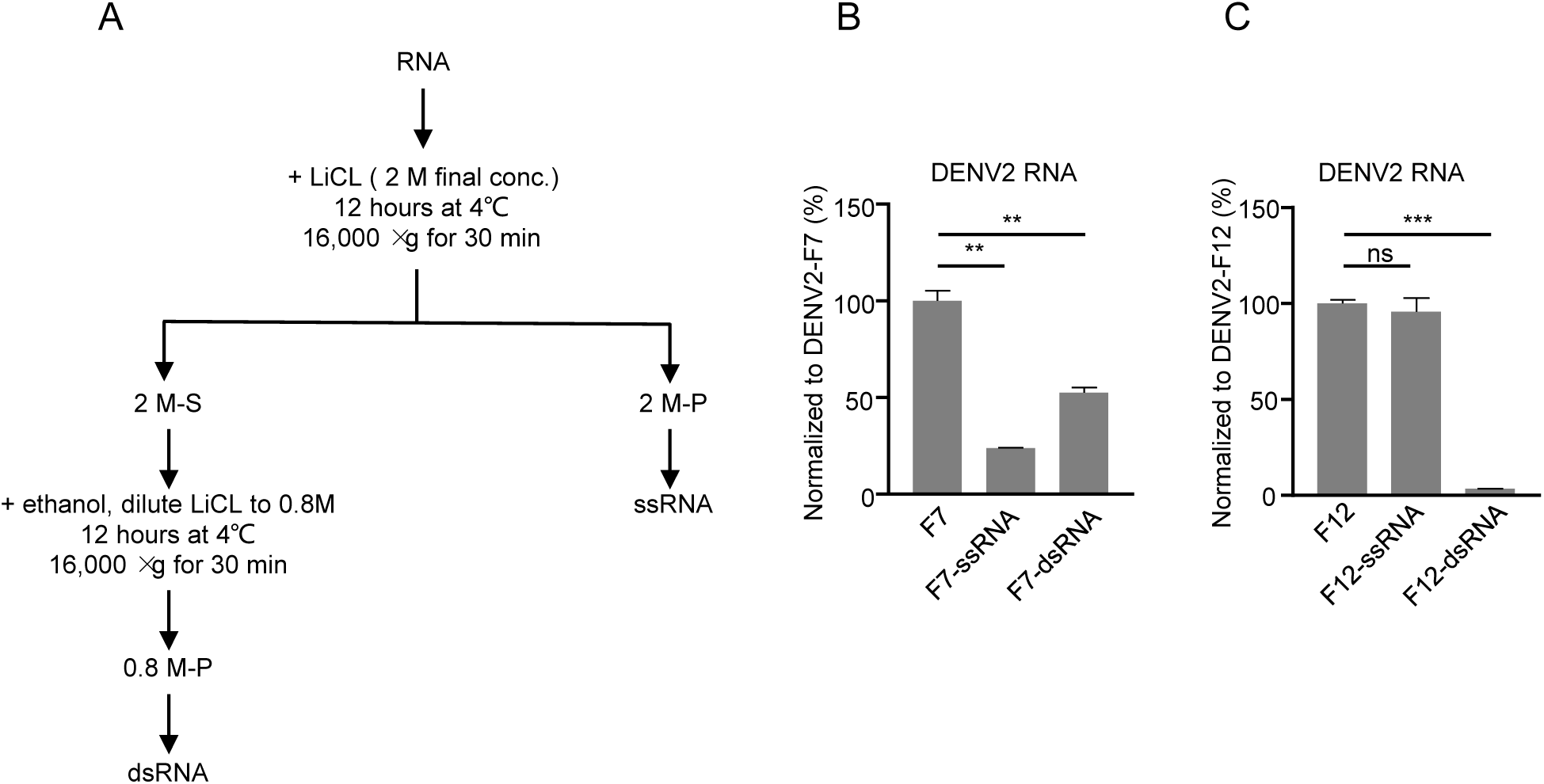
Purification of DENV2 PAMPs in the DENV2-F7 RNA through LiCl differential precipitation. (**A**) Schematic representation of separation of ssRNA and dsRNA through LiCl differential precipitation. LiCl was added to the RNA preparation to a final concentration of 2 M. ssRNA was precipitated and collected in pellet (2M-P) by centrifugation at 16,000 ×g for 30 min. The supernatant (2M-S) was added with ethanol to a final concentration of 60%, and the concentration of LiCl was decreased to 0.8M. dsRNA was precipitated and collected in pellet (0.8M-P) by centrifugation at 16,000 ×g for 30 min. (**B-C**) ssRNA and dsRNA in F7 (**B**) and F12 (**C**) derived from density gradient ultracentrifugation of DENV2-infected cellular RNA (see Fig S5) were separated by LiCl differential precipitation, and DENV2 RNA levels were analyzed by RT-qPCR, normalized to F7 or F12. The error bars represent standard deviations from two independent experiments. Student’s t test was used for statistical analysis. ns, P>0.05; **, p < 0.01; ***, p < 0.001.

**Fig S7.**
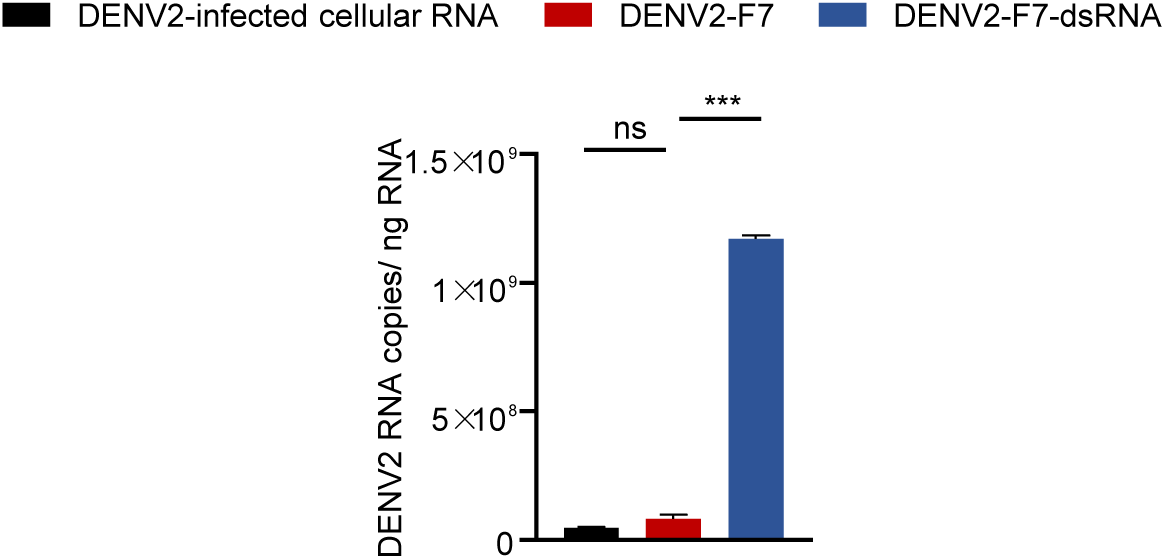
Viral RNA is enriched in the DENV2-F7-dsRNA preparation. Quantification of DENV2 RNA levels in DENV2-infected cellular RNA, DENV2-F7 and DENV2-F7-dsRNA. The error bars represent standard deviations from two independent experiments. Student’s t test was used for statistical analysis. ns, P>0.05; ***, p < 0.001.

**Fig S8.**
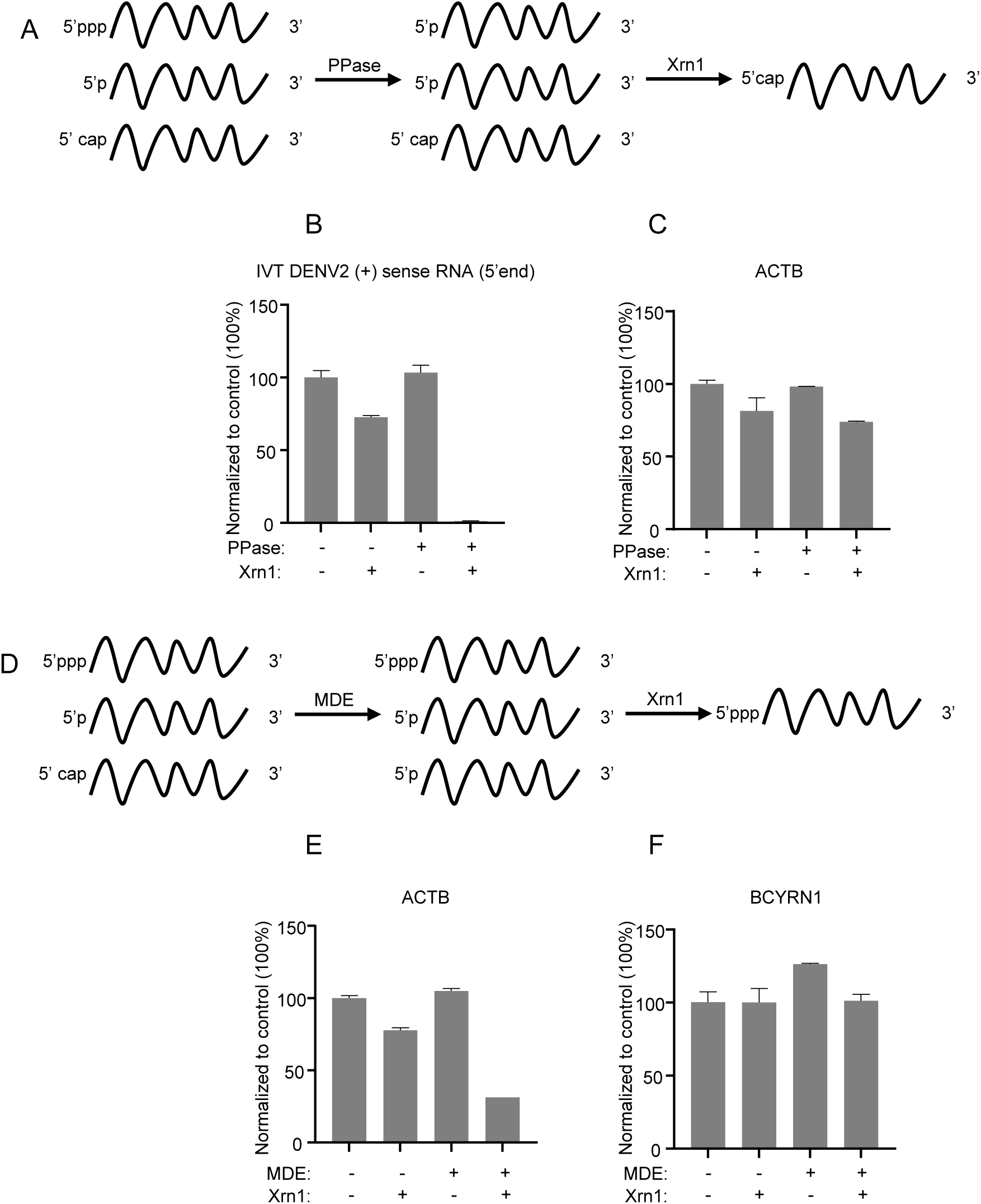
Analysis of the 5’-end modification of DENV2 PAMP RNA. (**A**) Schematic representation of the 5’ppp-specific RNA enzymatic digestion assay. RNA species are treated with PPase that removes the γ- and β-phosphates from the 5’ppp of an RNA to generate a 5’p end, and then digested with 5’–3’ exonuclease Xrn1, which specifically digests 5’p-RNA but not 5’ppp-RNA. RNA bearing a 5’ppp is converted to 5’p after the PPase digestion and becomes sensitive to the Xrn1 digestion. RNA with a 5’p, irrespective of the PPase treatment, is sensitive to the Xrn1 digestion. RNA with a 5’cap remains stable upon the PPase treatment and is resistant to the Xrn1 digestion. (**B-C**) *In vitro* transcribed DENV2 replicon (+) sense RNA was supplemented with uninfected Huh7 cellular RNA, subjected to treatment with Xrn1, PPase, or a combination of both PPase and Xrn1. The 5’end of DENV2 replicon (+) sense RNA (**B**) and ACTB mRNA (**C**) were detected by RT-qPCR. The RNA levels were normalized to the mock treatment control group. The error bars represent standard deviations from two independent experiments. (**D**) Schematic representation of the 5’cap-specific RNA enzymatic digestion assay. RNA species are treated with MDE that removes the cap-0 or cap-1 structure of an RNA to generate a 5’p end, and then digested with Xrn1. RNA with a 5’ppp remains stable upon MDE treatment and is resistant to the Xrn1 digestion. RNA with a 5’p, irrespective of MDE treatment, is sensitive to the Xrn1 digestion. RNA bearing a 5’cap is converted to 5’p after MDE digestion and becomes sensitive to the Xrn1 digestion. (**E-F**) Uninfected Huh7 cellular RNA was subjected to treatment with Xrn1, MDE, or a combination of both MDE and Xrn1. ACTB mRNA (**E**) and BCYRN1 RNA (**F**) were detected by RT-qPCR. The RNA levels were normalized to a control sample. The error bars represent standard deviations from two independent experiments.

**Fig S9.**
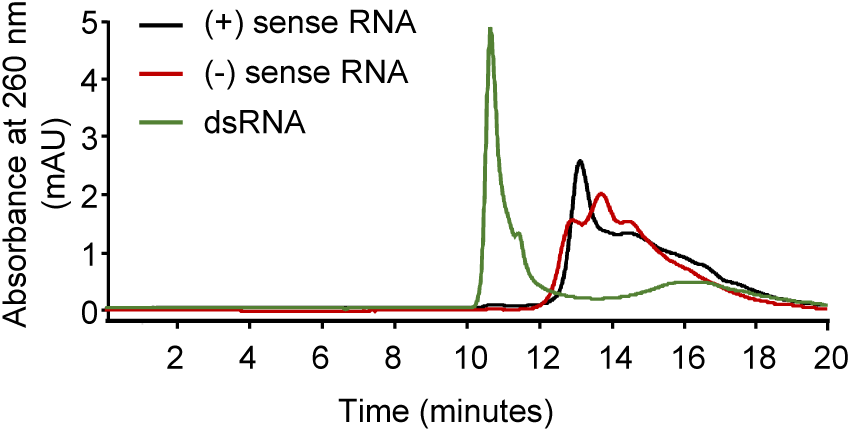
Analysis of single-stranded IVT DENV2 replicon (+) and (-) sense RNAs, as well as formed DENV2 RF dsRNA by size-exclusion chromatography. IVT DENV2 replicon (+) and (-) sense RNA, as well as DENV2 RF dsRNA were resolved on an Agilent Bio Size Exclusion Column SEC5, 2000 Å.

## References

1. Holmes EC, Twiddy SS. 2003. The origin, emergence and evolutionary genetics of dengue virus. Infect Genet Evol 3:19-28.

2. Islam MT, Quispe C, Herrera-Bravo J, Sarkar C, Sharma R, Garg N, Fredes LI, Martorell M, Alshehri MM, Sharifi-Rad J, Daştan SD, Calina D, Alsafi R, Alghamdi S, Batiha GE, Cruz-Martins N. 2021. Production, Transmission, Pathogenesis, and Control of Dengue Virus: A Literature-Based Undivided Perspective. Biomed Res Int 2021:4224816.

3. Shang W, Liu J, Yang J, Hu Z, Rao X. 2012. Dengue virus-like particles: construction and application. Appl Microbiol Biotechnol 94:39-46.

4. Klema VJ, Padmanabhan R, Choi KH. 2015. Flaviviral Replication Complex: Coordination between RNA Synthesis and 5’-RNA Capping. Viruses 7:4640-56.

5. Fernandez-Garcia MD, Mazzon M, Jacobs M, Amara A. 2009. Pathogenesis of flavivirus infections: using and abusing the host cell. Cell Host Microbe 5:318-28.

6. Guzman MG, Gubler DJ, Izquierdo A, Martinez E, Halstead SB. 2016. Dengue infection. Nat Rev Dis Primers 2:16055.

7. Akira S, Uematsu S, Takeuchi O. 2006. Pathogen recognition and innate immunity. Cell 124:783-801.

8. Hartmann G. 2017. Nucleic Acid Immunity. Adv Immunol 133:121-169.

9. Hei L, Zhong J. 2017. Laboratory of genetics and physiology 2 (LGP2) plays an essential role in hepatitis C virus infection-induced interferon responses. Hepatology 65:1478-1491.

10. Rehwinkel J, Gack MU. 2020. RIG-I-like receptors: their regulation and roles in RNA sensing. Nat Rev Immunol 20:537-551.

11. Hornung V, Ellegast J, Kim S, Brzózka K, Jung A, Kato H, Poeck H, Akira S, Conzelmann KK, Schlee M, Endres S, Hartmann G. 2006. 5’-Triphosphate RNA is the ligand for RIG-I. Science 314:994-7.

12. Pichlmair A, Schulz O, Tan CP, Näslund TI, Liljeström P, Weber F, Reis e Sousa C. 2006. RIG-I-mediated antiviral responses to single-stranded RNA bearing 5’-phosphates. Science 314:997-1001.

13. Schlee M, Roth A, Hornung V, Hagmann CA, Wimmenauer V, Barchet W, Coch C, Janke M, Mihailovic A, Wardle G, Juranek S, Kato H, Kawai T, Poeck H, Fitzgerald KA, Takeuchi O, Akira S, Tuschl T, Latz E, Ludwig J, Hartmann G. 2009. Recognition of 5’ triphosphate by RIG-I helicase requires short blunt double-stranded RNA as contained in panhandle of negative-strand virus. Immunity 31:25-34.

14. Schmidt A, Schwerd T, Hamm W, Hellmuth JC, Cui S, Wenzel M, Hoffmann FS, Michallet MC, Besch R, Hopfner KP, Endres S, Rothenfusser S. 2009. 5’-triphosphate RNA requires base-paired structures to activate antiviral signaling via RIG-I. Proc Natl Acad Sci U S A 106:12067-72.

15. Saito T, Owen DM, Jiang F, Marcotrigiano J, Gale M, Jr. 2008. Innate immunity induced by composition-dependent RIG-I recognition of hepatitis C virus RNA. Nature 454:523-7.

16. Kato H, Takeuchi O, Mikamo-Satoh E, Hirai R, Kawai T, Matsushita K, Hiiragi A, Dermody TS, Fujita T, Akira S. 2008. Length-dependent recognition of double-stranded ribonucleic acids by retinoic acid-inducible gene-I and melanoma differentiation-associated gene 5. J Exp Med 205:1601-10.

17. Ding Q, Huang B, Lu J, Liu YJ, Zhong J. 2012. Hepatitis C virus NS3/4A protease blocks IL-28 production. Eur J Immunol 42:2374-82.

18. Perry ST, Prestwood TR, Lada SM, Benedict CA, Shresta S. 2009. Cardif-mediated signaling controls the initial innate response to dengue virus in vivo. J Virol 83:8276-81.

19. Loo YM, Fornek J, Crochet N, Bajwa G, Perwitasari O, Martinez-Sobrido L, Akira S, Gill MA, García-Sastre A, Katze MG, Gale M, Jr. 2008. Distinct RIG-I and MDA5 signaling by RNA viruses in innate immunity. J Virol 82:335-45.

20. Sprokholt JK, Kaptein TM, van Hamme JL, Overmars RJ, Gringhuis SI, Geijtenbeek TBH. 2017. RIG-I-like Receptor Triggering by Dengue Virus Drives Dendritic Cell Immune Activation and T(H)1 Differentiation. J Immunol 198:4764-4771.

21. Sprokholt JK, Kaptein TM, van Hamme JL, Overmars RJ, Gringhuis SI, Geijtenbeek TBH. 2017. RIG-I-like receptor activation by dengue virus drives follicular T helper cell formation and antibody production. PLoS Pathog 13:e1006738.

22. Chazal M, Beauclair G, Gracias S, Najburg V, Simon-Lorière E, Tangy F, Komarova AV, Jouvenet N. 2018. RIG-I Recognizes the 5’ Region of Dengue and Zika Virus Genomes. Cell Rep 24:320-328.

23. Chang TH, Liao CL, Lin YL. 2006. Flavivirus induces interferon-beta gene expression through a pathway involving RIG-I-dependent IRF-3 and PI3K-dependent NF-kappaB activation. Microbes Infect 8:157-71.

24. da Conceição TM, Rust NM, Berbel AC, Martins NB, do Nascimento Santos CA, Da Poian AT, de Arruda LB. 2013. Essential role of RIG-I in the activation of endothelial cells by dengue virus. Virology 435:281-92.

25. Nasirudeen AM, Wong HH, Thien P, Xu S, Lam KP, Liu DX. 2011. RIG-I, MDA5 and TLR3 synergistically play an important role in restriction of dengue virus infection. PLoS Negl Trop Dis 5:e926.

26. Castillo Ramirez JA, Urcuqui-Inchima S. 2015. Dengue Virus Control of Type I IFN Responses: A History of Manipulation and Control. J Interferon Cytokine Res 35:421-30.

27. Coldbeck-Shackley RC, Eyre NS, Beard MR. 2020. The Molecular Interactions of ZIKV and DENV with the Type-I IFN Response. Vaccines (Basel) 8.

28. Gack MU, Diamond MS. 2016. Innate immune escape by Dengue and West Nile viruses. Curr Opin Virol 20:119-128.

29. Lee MF, Voon GZ, Lim HX, Chua ML, Poh CL. 2022. Innate and adaptive immune evasion by dengue virus. Front Cell Infect Microbiol 12:1004608.

30. Morrison J, Aguirre S, Fernandez-Sesma A. 2012. Innate immunity evasion by Dengue virus. Viruses 4:397-413.

31. Uno N, Ross TM. 2018. Dengue virus and the host innate immune response. Emerg Microbes Infect 7:167.

32. Shalem O, Sanjana NE, Hartenian E, Shi X, Scott DA, Mikkelson T, Heckl D, Ebert BL, Root DE, Doench JG, Zhang F. 2014. Genome-scale CRISPR-Cas9 knockout screening in human cells. Science 343:84-87.

33. Uzri D, Gehrke L. 2009. Nucleotide sequences and modifications that determine RIG-I/RNA binding and signaling activities. J Virol 83:4174-84.

34. Xu Y, Zhong J. 2016. Innate immunity against hepatitis C virus. Curr Opin Immunol 42:98-104.

35. Feng Q, Hato SV, Langereis MA, Zoll J, Virgen-Slane R, Peisley A, Hur S, Semler BL, van Rij RP, van Kuppeveld FJ. 2012. MDA5 detects the double-stranded RNA replicative form in picornavirus-infected cells. Cell Rep 2:1187-96.

36. Rehwinkel J, Tan CP, Goubau D, Schulz O, Pichlmair A, Bier K, Robb N, Vreede F, Barclay W, Fodor E, Reis e Sousa C. 2010. RIG-I detects viral genomic RNA during negative-strand RNA virus infection. Cell 140:397-408.

37. Holmes EC. 1998. Molecular epidemiology and evolution of emerging infectious diseases. Br Med Bull 54:533-43.

38. Limthongkul J, Mapratiep N, Apichirapokey S, Suksatu A, Midoeng P, Ubol S. 2019. Insect anionic septapeptides suppress DENV replication by activating antiviral cytokines and miRNAs in primary human monocytes. Antiviral Res 168:1-8.

39. Cleaves GR, Ryan TE, Schlesinger RW. 1981. Identification and characterization of type 2 dengue virus replicative intermediate and replicative form RNAs. Virology 111:73-83.

40. Schlee M. 2013. Master sensors of pathogenic RNA - RIG-I like receptors. Immunobiology 218:1322-35.

41. Choi KH. 2021. The Role of the Stem-Loop A RNA Promoter in Flavivirus Replication. Viruses 13.

42. Wengler G, Wengler G, Gross HJ. 1978. Studies on virus-specific nucleic acids synthesized in vertebrate and mosquito cells infected with flaviviruses. Virology 89:423-37.

43. Sherwood AV, Rivera-Rangel LR, Ryberg LA, Larsen HS, Anker KM, Costa R, Vågbø CB, Jakljevič E, Pham LV, Fernandez-Antunez C, Indrisiunaite G, Podolska-Charlery A, Grothen JER, Langvad NW, Fossat N, Offersgaard A, Al-Chaer A, Nielsen L, Kuśnierczyk A, Sølund C, Weis N, Gottwein JM, Holmbeck K, Bottaro S, Ramirez S, Bukh J, Scheel TKH, Vinther J. 2023. Hepatitis C virus RNA is 5’-capped with flavin adenine dinucleotide. Nature 619:811-818.

44. Vabret N, Najburg V, Solovyov A, Gopal R, McClain C, Šulc P, Balan S, Rahou Y, Beauclair G, Chazal M, Varet H, Legendre R, Sismeiro O, Sanchez David RY, Chauveau L, Jouvenet N, Markowitz M, van der Werf S, Schwartz O, Tangy F, Bhardwaj N, Greenbaum BD, Komarova AV. 2022. Y RNAs are conserved endogenous RIG-I ligands across RNA virus infection and are targeted by HIV-1. iScience 25:104599.

45. Paul AV, Wimmer E. 2015. Initiation of protein-primed picornavirus RNA synthesis. Virus Res 206:12-26.

46. Chiang JJ, Sparrer KMJ, van Gent M, Lässig C, Huang T, Osterrieder N, Hopfner KP, Gack MU. 2018. Viral unmasking of cellular 5S rRNA pseudogene transcripts induces RIG-I-mediated immunity. Nat Immunol 19:53-62.

47. Runge S, Sparrer KM, Lässig C, Hembach K, Baum A, García-Sastre A, Söding J, Conzelmann KK, Hopfner KP. 2014. In vivo ligands of MDA5 and RIG-I in measles virus-infected cells. PLoS Pathog 10:e1004081.

48. Sanchez David RY, Combredet C, Sismeiro O, Dillies MA, Jagla B, Coppée JY, Mura M, Guerbois Galla M, Despres P, Tangy F, Komarova AV. 2016. Comparative analysis of viral RNA signatures on different RIG-I-like receptors. Elife 5:e11275.

49. Zhao Y, Ye X, Dunker W, Song Y, Karijolich J. 2018. RIG-I like receptor sensing of host RNAs facilitates the cell-intrinsic immune response to KSHV infection. Nat Commun 9:4841.

50. Srikiatkhachorn A, Mathew A, Rothman AL. 2017. Immune-mediated cytokine storm and its role in severe dengue. Semin Immunopathol 39:563-574.

51. Li L, Gan T, Ma Z, Huang Y, Zhong J. 2024. Assessing risk of Bombali virus spillover to humans by mutagenesis analysis of viral glycoprotein. hLife 2:32-42.

52. He Z, Ye S, Xing Y, Jiu Y, Zhong J. 2021. UNC93B1 curbs cytosolic DNA signaling by promoting STING degradation. Eur J Immunol 51:1672-1685.

53. Xie X, Zou J, Zhang X, Zhou Y, Routh AL, Kang C, Popov VL, Chen X, Wang QY, Dong H, Shi PY. 2019. Dengue NS2A Protein Orchestrates Virus Assembly. Cell Host Microbe 26:606-622.e8.

